# Ancient DNA Reveals the Lost Domestication History of South American Camelids in Northern Chile and Across the Andes

**DOI:** 10.1101/2020.10.16.337428

**Authors:** Paloma Díaz-Maroto, Alba Rey-Iglesia, Isabel Cartajena, Lautaro Núñez, Michael V Westbury, Valeria Varas, Mauricio Moraga, Paula F. Campos, Pablo Orozco-terWengel, Juan Carlos Marín, Anders J. Hansen

## Abstract

The study of South American camelids and their domestication is a highly debated topic in zooarchaeology. Identifying the domestic species (alpaca and llama) in archaeological sites based solely on morphological data is challenging due to their similarity with respect to their wild ancestors. Using genetic methods also present challenges due to the hybridization history of the domestic species, which are thought to have extensively hybridized following the Spanish conquest of South America that resulted in camelids slaughtered en-masse. In this study we generated mitochondrial genomes for 61 ancient South American camelids dated between 3,500 - 2,400 years before the present (Early Formative period) from two archaeological sites in Northern Chile (Tulán 54 and 85), as well as 66 modern camelid mitogenomes and 815 extant mitochondrial control region sequences from across South America. In addition, we performed osteometric analyses to differentiate big and small body size camelids. A comparative analysis of these data suggests that a substantial proportion of the ancient vicuña genetic variation has been lost since the Early Formative period as it is not present in modern specimens. Moreover, we propose a model of domestication that includes an ancient guanaco population that no longer exists. Finally, we find evidence that interbreeding practices were widespread during the domestication process by the early camelid herders in the Atacama during the Early Formative period and predating the Spanish conquest.

## Introduction

The study of South American camelids (SACs) and their domestication has been a central subject in South American zooarchaeology since the 1970’s. Many studies have focused on elucidating the domestication process as well as use and exploitation of camelid species in the past. These animals were extremely plastic, able to adapt to a great variety of environments found across the Andes, and were even present in remote islands, emphasizing their importance in local ecosystems and for historic/prehistoric human populations. Camelids were key in the transition from hunter-gatherer to a mixed economy and played a central role in the cosmic vision of past Andean communities. (**Wing, 1972**; **Stahl, 1988**; **Stanley et al., 1994**; **Wheeler, 1995; Olivera, 1997; Yacobaccio et al., 1998**; **Cartajena, 2002; Izeta et al., 2009**; **L’Heureux, 2010**; **Yacobaccio et al., 2013**; **Gasco et al., 2014).**

There are currently four camelid species inhabiting South America, two wild species, guanaco (*Lama guanicoe*) and vicuña (*Vicugna vicugna*); and two domestic species, llama (*Lama glama*) and alpaca (*Vicugna pacos*). Guanaco and vicuña are the most important endemic large herbivores in South America due to their ecological dominance in the upper Andean ecosystem and their importance for human populations (**Muñoz & Mondini, 2008**). Llama and alpaca first appeared in the fossil record around 7,000 years before present (yrBP) as the result of domestication carried out by human communities across the Andes (Wheeler, Pires-Ferreira, & Kaulicke, 1976; Wheeler, 1984; Wheeler, 1995).

Several different hypotheses have been proposed for the origin of the domestic species (reviewed in **Wheeler, 1995**): 1) llamas were domesticated from guanacos and alpacas were domesticated from vicuñas, 2) llamas were domesticated from guanacos and alpacas derived from hybridization between llamas and vicuñas, 3) llamas and alpacas were both domesticated from guanacos while vicuña was never domesticated, and 4) llama and alpaca evolved from extinct wild precursors, while guanaco and vicuña were never domesticated.

To elucidate the origin and evolutionary history of camelids, numerous studies have used DNA analyses of extant specimens. However, the conclusions drawn from these studies are ambiguous. Some mitochondrial DNA results supporting that domestication of guanacos giving rise to both domestic species. Others support the concomitant domestication of guanacos and vicuñas into llamas and alpacas. Finally, efforts using whole genome sequencing suggest support for the hypothesis that llama derived from guanaco and alpaca derived from vicuña (**Stanley et al., 1994; Kadwell et al., 2001; Wheeler et al.,2006; Marín et al., 2007a, Fan et al., 2020**). Although some of these problems may be addressed using ancient DNA samples, such material is scarce for SACs with only four complete mitochondrial genomes from two different species currently available, three llamas dated to around 700 yr BP from Isla Mocha in Southern Chile (**Westbury et al., 2016**) and one vicuña dated between 3200-2400 yr BP, from Tulán 54 (**Diaz-Maroto et al., 2018**).

Osteometry has also been widely used in studies trying to understand the domestication of SACs. The first phalanx (anterior and posterior) and astragalus (**Yacobaccio, 2003; Gallardo & Yacobaccio, 2005**; **Izeta & Cortés, 2006**; **Cartajena, Núñez, & Grosjean, 2007; Cartajena, 2009; Izeta et al., 2009; L’Heureux, 2010; Reigadas, 2012, 2014; Gasco et al., 2014**) have been used for classification of two distinct size groups, the large sized guanaco and llama (genus *Lama*) and the small sized vicuña and alpaca (genus *Vicugna*; **Wheeler, 1995**). However, due to the lack of significant differences between the domestic and wild camelids within each group (Cartajena et al., 2007); the taxonomic assignment within each genus remains challenging.

To evaluate the presence of domestic camelids in the Early Formative period in Northern Chile and to elucidate the origin and evolutionary history of South American camelids, we use a combination of osteometry and mitogenomics of ancient samples collected in the archeological site of Tulán in northern Chile (Figure 1) and modern samples collected throughout the South American distribution of these species. The comparison between the patterns of genetic variation of this dataset of ancient (60) and modern (66) mitogenomes along with the largest dataset of SAC mitochondrial D-Loop sequences (815) provides a unique and deeper insight into one of the biggest unsolved mysteries of South American archaeozoology, the South American Camelid domestication.

**Figure 1.**
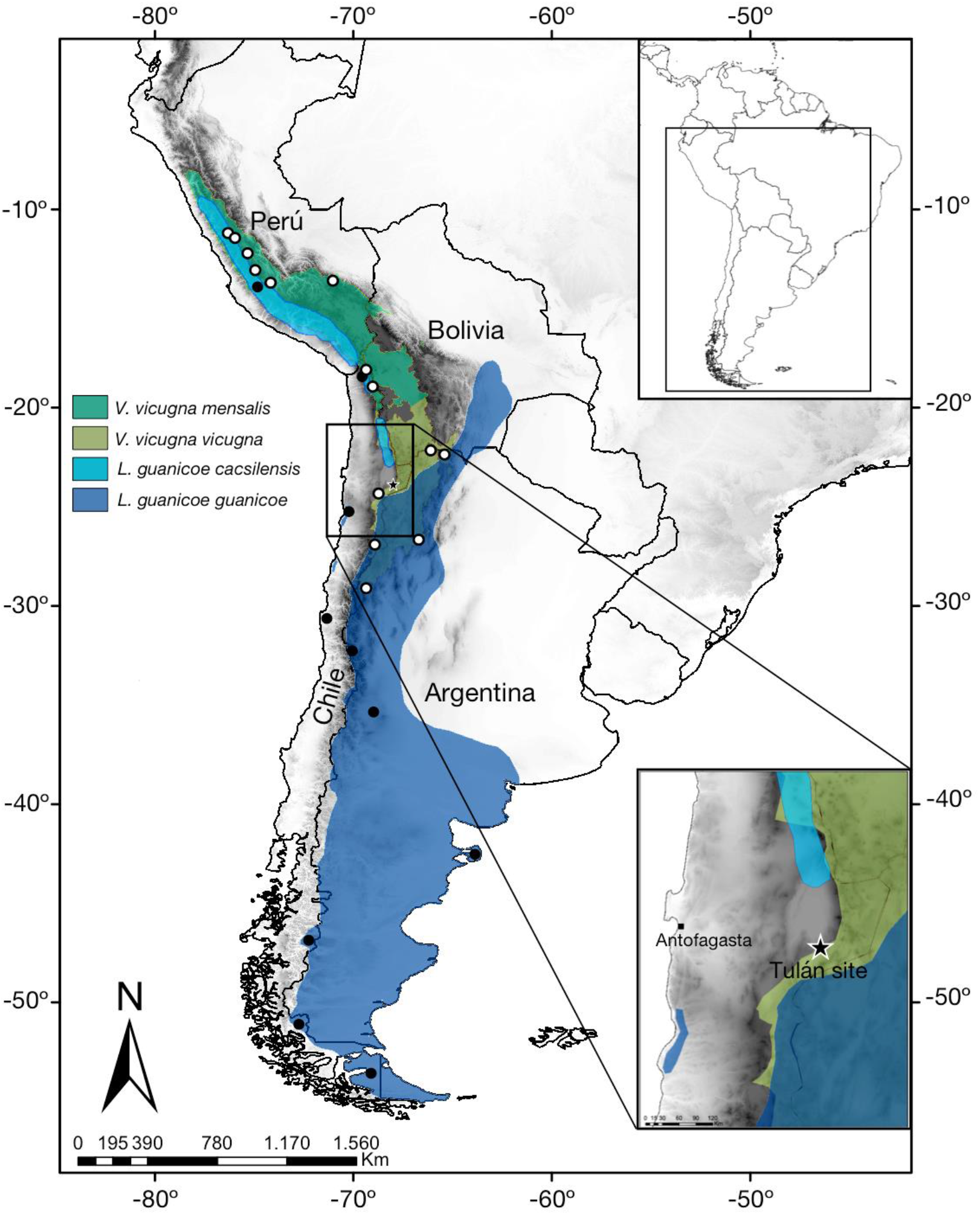
Distribution map of two guanaco and vicuña subspecies. Black and white dots represent the sampling locations of modern guanacos and modern vicuñas respectively. A star shows the Tulán site.

## Materials and methods

### Archeological samples

A total of 75 specimens from putative camelid remains, collected in several archaeological excavations in the North of Chile between 2003 and 2009 (**Cartajena, 2003, 2006, 2009; Cartajena & López, 2011**), were selected for DNA analyses (**Table S1**). These samples were collected in the archaeological sites Tulán-54 (3,080 - 2,380 yr BP; 57 samples) and Tulán-85 (3,140 - 2,660 yr BP; 18 samples), belonging to the Early Formative period. Samples from Tulán-54 were selected from inside and outside of the ceremonial center (35 phalanges and 22 astragalus) in order to cover a wide sampling range because of different use of the space in the site. In Tulán-85, samples were collected from an inhabited area (16 phalanges and 2 astragalus) (See Supplementary material for more information about the principal sites). We additionally collected two samples from the sites Tulán-52 and Tulán-94 to gain more information into the camelid domestication process in this area as a whole. Tulán-52 is one of the most important Late Archaic sites in Northern Chile and is dated to between 4,340 and 4,270 yr BP, (**Aldenderfer, 1989; Núñez et al., 2006**). Tulán-94 is another residential site located along the Tulán ravine dating from 2,890 - 3,640 yr BP (Early Formative period). The stratified deposits have recorded skeletal remains, mainly camelid remains as well as plants, shells, and beads among others (Núñez et al., 2006).

### Morphometric analysis

We recorded osteometric measurements from 61 out of the 75 bone samples from the archaeological sites using Vernier calipers to the nearest 0.1 mm, as described by von den Driesch (**von den Driesch, 1976**). Due to fragmentation or bad preservation, 18 out of 61 samples were not used for morphometric analysis. Osteometric measures were performed on each bone sample prior to bone drilling for aDNA to avoid bias due to damage during drilling. Measurements were performed on phalanges and astragalus. Bones were selected from different levels and squares in the excavation sites in order to choose different individuals (**Table S4**). Only adult individuals (determined by a phalanx with fused epiphysis and astragalus) were selected for the morphometric analysis. The first phalanx was selected due to its high frequency in the sites and because it is easy to differentiate between anterior and posterior phalanges. Breadth of proximal articulation (BFp) and the depth of the proximal epiphysis (Dp) measures were taken from the proximal side of the phalanx. Astragalus measures included the breadth distal (Bd) and greatest length medial (GLm) (**von den Driesch 1976**). Morphological differentiation between “small size animals” and “big size animals” was tested using correlations between the anterior and posterior phalanges and astragalus using the program PAST (Hammer et al.,2001). In order to increase the sample size and robustness of our analyses, our data was combined with published morphological data from the same sites (**Cartajena, 2003; 2006; 2009**) and measures from the anterior phalanx for the four modern camelid species (**L’Hereux, 2010**).

### Ancient DNA extraction

DNA was extracted from 61 samples from phalanx and astragalus bones in a dedicated ancient DNA laboratory at the Centre for Geogenetics, University of Copenhagen. From each specimen, 150 mg of bone was drilled using a Dremel drill to obtain bone powder for DNA extraction. DNA extractions were performed as in **Dabney et al., (2013)** with the following modifications: samples were incubated overnight with the extraction buffer at 42 °C instead of at 37 °C, the bone powder was pelleted out of suspension, and the supernatant concentrated down to 150 - 200 μl for each sample using 30 kDa Amicon centrifugal filter unit (Milipore). Binding buffer 13 times larger in volume was added to the concentrated supernatant and DNA was purified with MinElute columns (Qiagen) following the manufacturer’s instructions with the exception of a 15-minute incubation at 37 °C during the elution step. DNA was eluted by adding 50 μl of EB buffer twice for a total volume of 100 μl, in order to increase DNA yields. For every round of extractions, one blank extraction control was used to monitor for contamination.

### Ancient DNA Library build

Double stranded libraries were built using 21.25 μl of the extracts following the protocol from Meyer & Kircher (2010) with some modifications: the initial DNA fragmentation was not performed and MinElute kit (Qiagen) was used for the purification steps. A quantitative PCR (qPCR) assay was performed prior to the amplification step to assess the optimal number of cycles for index PCR to avoid overamplification using Roche LC480 SYBR Green I Master Mix with the primers IS5 and IS6 from **Meyer & Kircher (2010)**. All qPCR were carried out in a total volume of 20 μl with the following reaction conditions: 95 °C/10min, 35 cycles of 95 °C/30 sec, 55 °C/30 sec, 72 °C/30 sec followed by 1 cycle at 95 °C/30 sec, 55 °C/1 min and a detection final step at 95 °C/30 sec. Indexing PCRs were performed in 50 μl reactions using 25 μl KAPA HiFi HotStart Uracil+ReadyMix, 2 μl of forward InPE 1.0 (10 μM) (Meyer & Kircher, 2010), 2 μl of an index primer containing a unique 6-nucleotide index tag (10 Mm), and 21 μl of DNA library. The cycling conditions of amplification PCR were 45sec at 98 °C, 14-17 cycles of (98 °C 15 sec, 65 °C 30 sec, 72 °C 30 sec) and a final elongation at 72 °C for 1 min. MinElute columns were used for purification after index PCR and PCR products were quantified using a Qubit Fluorometer (HS). In order to reach the required 300 - 500 ng of starting material for performing the capture experiments, some of the libraries were reamplified. A reamplification PCR was setup with 10.5 μl of the indexed library, 12.5 μl of KAPA HiFi HotStart Uracil+ReadyMix and 1 μl of each re-amplification primer, IS5 and IS6 (10 μM stock concentration) (Meyer & Kircher, 2010). Cycling conditions were the same as for the index PCR but 7 cycles for annealing. PCR products were purified and quantified as indicated above.

### Ancient DNA Mitochondrial capture

Our sequencing libraries underwent target-capture prior to sequencing in order to enrich our libraries of endogenous mitochondrial DNA sequences. We designed a camelid set bait which was composed of 120 nucleotides baits tiled every four bases across a representative sample of four living camelid mitochondrial genome sequences. The 6,234 baits were then synthesized at MYcroarray (https://arborbiosci.com/) as part of several MYbaits targeted enrichment kits. Hybridization by capture was performed following the MYbaits v3 manual, using 350 - 500 ng of library starting material. We performed a single round of enrichment at 55 °C for 24 hours. Enriched libraries were eluted in a total volume of 30 μl. Post-capture amplification was carried out according to the suggested protocol using KAPA HiFi HotStart Uracil+ReadyMix and 12 cycles for the cycling PCR conditions. Enriched libraries were quantified using Agilent 2200 TapeStation System, pooled in equimolar amounts, and sequenced on an Illumina HiSeq 2500 using 80bp single-end chemistry at the Danish National High-Throughput Sequencing Center.

### Ancient DNA bioinformatic processing

PALEOMIX was used to perform basic read processing **(Schubert et al., 2014):** (i) adapter trimming, (ii) mapping trimmed reads to the reference mitogenome of *Lama pacos* (GenBank: Y19184.1) and (iii) PCR duplicate removal. Seeding was disabled for mapping and low-quality reads of less than 30 bp were discarded. The BAM files with unique reads were loaded into Geneious v7 **(Kearse et al., 2012)** to generate mitochondrial consensus sequences for each sample. Mitochondrial consensus sequences were generated using the “strict” consensus that calls bases using majority rule and requires a minimum depth-of-coverage of 3x **(Table S5)**. This resulted in the generation of 61 complete mitochondrial genomes. DNA damage and fragmentation patterns were also estimated within the PALEOMIX pipeline using MapDamage2 (**Jónsson et al., 2013**) (Figure S5).

### Data set of new modern DNA samples

For comparison with the aDNA samples, blood and tissue samples were obtained from 66 georeferenced modern samples to generate the whole mitochondrial genome excluding the Control Region (15438bp; Figure 1, Table S3). These include 16 guanacos: Northern guanaco (6 *L. guanicoe cacsilensis*) and Southern guanaco (10 *L. guanicoe guanicoe*); 17 vicuñas: Northern vicuña (6 *V. vicugna mensalis*) and Southern vicuña (11 *V. vicugna vicugna*) from throughout the current distribution across Peru, Argentina and Chile; 16 llamas from a slaughterhouse in Putre, Chile; and 17 alpacas from a farmer and breeder in Chile. Genomic DNA was extracted using the Puregene Tissue Core Kit A (Qiagen). For each individual, 1–3 μg of DNA was sheared into fragments of 150–700 bp prior to the construction of the sequencing library. Adapter ligation (or library build) was performed using the NNNPaired-end sequencing library kit with an insert size of approximately 300 bp. Libraries were sequenced to mid/low-coverage (11-14x) on an Illumina Hiseq X Ten platform.

From the resultant raw reads, we removed reads with ≥ 10% unidentified nucleotides, >10 nucleotides aligned to the adaptor, with ≤10% mismatches allowed, with >50% bases having phred quality <5 and putative PCR duplicates generated during library construction. Following the initial processing, unpaired reads were removed using Trimmomatic **(Bolger, Lohse & Usadel, 2014)**, and reads with a Phred quality score lower than 33. High quality reads were aligned to conspecific reference mitochondrial genomes (access number NCBI, EU681954; FJ456892; AJ566364; AP003426) as appropriate using BWA **(Li, 2009, 2013)**, the resultant Bayesian alignments were sorted and filtered using SAMtools program (**Li, 2009)** to sort the reads and filter them. The latter commands included filters to eliminate optical duplicates, unmapped reads and mates, and only keeping reads that mapped completely. We reconstructed the mitochondrial consensus sequences using GenomeAnalysisTKjar from GATK with references indexed with Picard **(Li, 2014)**.

### Mitochondrial sequences compilation

We included 300 base pair (bp) mitochondrial control region sequences from 815 modern camelids representing the Northern guanaco (54 *L. guanicoe cacsilensis*), the Southern guanaco (366 *L. guanicoe guanicoe*), the Northern vicuña (218 *V. vicugna mensalis*), the Southern vicuña (35 *V. vicugna vicugna*), domestic alpaca (53 *V. pacos*), domestic llama (79 *L. glama*) and ten hybrids (three hybrids between the two vicuña forms and seven hybrids between llama and guanaco; **Table S2**). Additionally, three ancient guanaco mitogenomes from Isla Mocha (Chile) (KX388532.1; KX388533.1; KX388534.1) one guanaco (NC_011822.1), one vicuña mitogenome (FJ456892.1), one llama mitogenome (AP003426.1) and two alpaca mitogenomes (KU168760.1; AJ566364.1) were downloaded from NCBI and included in the analyses (Table S3 and Figure S4).

### Phylogenetic and Demographic Analysis using mitogenome data

The 131 sequences from ancient and modern South American camelid mitogenomes and one Bactrian camel sequence used as outgroup, were aligned using Geneious V7 resulting in an alignment of 15,456 bp, including protein-coding genes, transfer RNA (tRNA), ribosomal RNA (rRNA) genes, and a partial control region sequence (CR).

Partition Finder v1.1.1. (**Lanfear et al., 2012**) was used to determine the optimal partitioning scheme for this alignment and substitution model for downstream analyses. Several partition schemes were tested **(Table S6)**. The best fitting-model to our alignment consisted of five partitions (tRNA and rRNA as one partition each and protein-coding genes divided by codon as three separate partitions) with HKY+I+G for partitions 1, 2, 3 and 5 and GTR+I+G for partition 4 as substitution model.

Phylogenetic relationships were estimated using the software MrBayes v3.2.6 (**Ronquist & Huelsenbeck, 2003**) and the substitution evolution models identified above. The MrBayes MCMC algorithm was run twice with three cold and one hot chain for 1×10^6^ generations, sampling every 1×10^3^ generations and discarding the first 25% of steps of the MCMC as burn-in following visual examination of the stabilization of the likelihood values of the inferred tree. Trees were summarized with the majority-rule consensus approach, using posterior probability as a measure of clade support. The final consensus tree was visualized in FigTree v.1.4.2 **(Rambaut, 2014).** The number of distinct haplotypes, haplotype diversity and nucleotide diversity were estimated in R using pegas v0.12 (Paradis 2010) and ape v5.3 (Paradis & Schliep 2019). We estimated nucleotide diversity excluding sites with gaps and missing data in each pairwise comparison.

Additionally, ultrametric trees were estimated for the dataset using BEAST v.1.8.0 (**Drummond et al 2012)** under a coalescent model and with *C. dromedarius* and *C. bactrianus* as outgroups. A Shimodaira-Hasegawa test for the evolution of the trio vicugna, guanaco and dromedary was calculated in MEGA X (Molecular Evolutionary Genetics Analysis across computing platforms) (Kumar et al., 2018) and showed no significant evidence of deviation of the data from the molecular clock null hypothesis. Thus, the number of observed substitution differences between the pair vicugna/dromedary (guanaco/dromedary had the same value) were used to estimate a substitution rate of 2.28-e8 per million years (based on timetree.org (Kumar et al 2017) divergence of 20.56 million years reported between these two taxa) for the BEAST analysis. A total of 20 million steps of BEAST’s MCMC algorithm were carried out discarding the first 10% as burn-in. Stability of the BEAST run was determined by achieving ESS values larger than 200 which were visualised in Tracer (Rambaut & Drummond 2007) and convergence was determined by running the analysis in duplicate and obtaining the same tree. To investigate changes of population size through time, a Bayesian Skyline reconstruction was performed on the data using BEAST and visualised in Tracer v1.4.1 (Rambaut & Drummond 2007). The plot was constructed with a rate of base substitution of 1,2% substitution per base pair per million years and HKY as a model of nucleotide substitution with a MCMC of 10 million steps following a discarded first 10% of each chain as a burn-in.

### Haplotype networks using mitochondrial control regions sequences

To explore the relationship between the different subspecies and their wild relatives, we aligned the partial CR (control region) from the ancient and modern mitogenomes with a total of 849 specimens for the four species covering the distribution range across Peru, Chile, Argentina and Ecuador using MAFFT (Katoh & Standley, 2013) **(Table S2)**. The final alignment consisted of 300 bp from the hypervariable I domain. We used this dataset to calculate a haplotype median-joining network using PopART **(Leigh et al., 2015)** and to investigate the change in haplotype composition over time using TempNet (**Prost & Anderson, 2011**) by dividing the dataset into two time periods, i.e. the 41 ancient samples and the 808 modern samples.

## Results

### Morphometric analysis of the phalanx and astragalus

Morphometric analysis of the astragalus and anterior and posterior phalanges show two differentiated groups based on body size, small sized and big sized individuals (Figure 2). The cutting point for BFp was 18mm to separate between small and big size camelids (Cartajena et al, 2007; Cartajena, 2009). The small size cluster most likely represents alpaca and vicuña specimens, while guanaco and llama individuals most likely form the big size cluster. Modern camelids are shown in red font in graphic with standard deviations (L’Hereux, 2010) (Figure 2).

**Figure 2.**
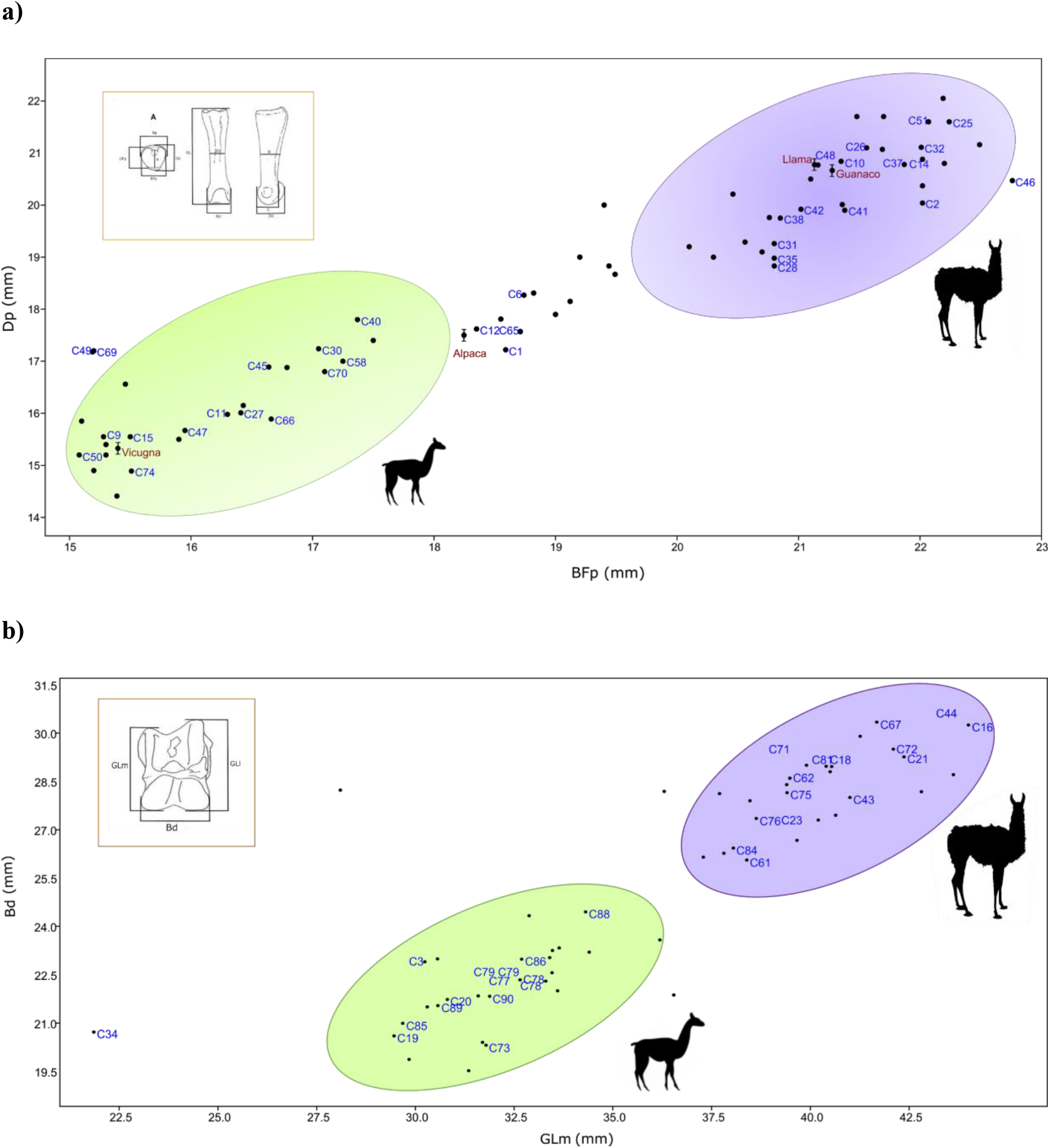
Correlation graphics of anterior phalanx and astragalus measurements. (a) Correlation for the anterior phalanx measurements; Dp = Depth proximal and Bfp = Breath of Facies articularis proximalis. (b) Correlation for the astragalus measurements; Bd = Breadth distal and GLm = Greatest length medial. Ancient camelid samples sequenced are shown in blue font and the black points correspond to other samples from the same archaeological site but for which were not included in DNA analyses. Modern camelids are shown in red font in graphic with standard deviations (L?ereux, 2010) (a). The green ellipsis indicates the group of small sized specimens and the purple ellipsis indicates the group of large sized specimens.

### Ancient mitogenome sequencing

Using a capture strategy to enrich our aDNA libraries for mitochondrial sequences we generated 61 near complete mitogenome sequences from camelid bone samples collected in the Tulán ravine and Atacama Salar: 57 from Tulán-54, 18 from Tulán-85, and two from Tulán-94 and Tulán-52. Average depth of coverage in the assembled mitogenomes ranged between 2.53x and 153.03x with a final length of 15,438 bp.

The analysis with mapDamage showed the characteristic pattern of ancient DNA consistent with postmortem deamination damage in all aDNA samples (Figure S5). An increasing presence of CT and G-A mutations at the terminal ends of sequenced molecules were detected at the 5’-ends and 3’termini respectively, as expected from the damage model used, generating an excess of cytosine deaminations at single-strands ends of the DNA templates.

### Phylogenetic analysis

The Bayesian phylogenetic analysis of the ancient and modern mitogenomes using the Bactrian camel as an outgroup resulted in strong statistical support for four clades in our data (Figure 3). Clades 1, 2 and 3 form a large monophyletic clade supported by a posterior probability of 1 with all modern domestic camelids located in Clades 1 and 3. Clade 1 consists of a mixture of modern llama and alpaca mitogenomes and 20 ancient samples (three from Tulán-85, 16 from Tulán-54 and one from Tulán-52) with 13 individuals morphologically identified as large sized animals and one specimen as intermediate sized. Most modern guanacos grouped within Clade 2, together with nine samples from Tulán-54 morphologically identified as big and intermediate sized animals, while Clade 3 consists of both domestic camelid species and includes one modern guanaco, as well as five ancient samples (C44, C5, C77, C52 and C48) that are closer to the ancestral mitogenome of the big clade. Clade 4 was supported by a posterior probability of one and grouped the majority of modern wild *V. vicugna* and two alpacas (*V. pacos*) (A_1557_alpaca and A03_alpaca) together with 15 ancient specimens (12 from Tulán-54 and 3 from Tulán-85) that were morphologically identified as small size camelids. The haplotype diversity for the ancient samples was 1.00 and for the modern samples, the values oscillated within 0,971 to 0,993 (Table S9).

**Figure 3.**
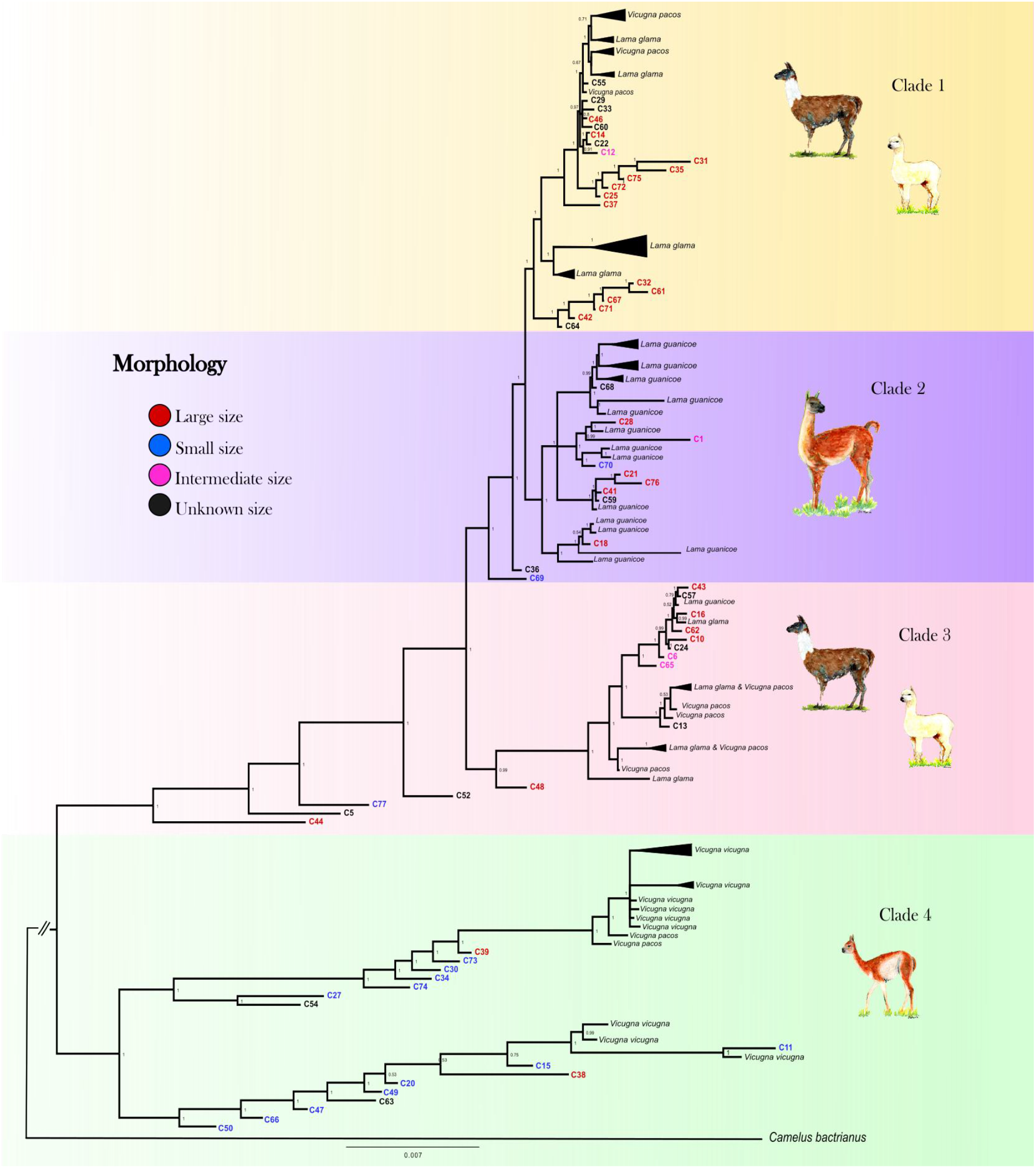
Mitogenomic phylogenetic tree. Bayesian reconstruction of phylogenetic relationships between ancient and modern camelids using complete mitochondrial genome data. For the aDNA samples their identification code appears on the branch tip with a colour indicative of their size class assignment based on the sample’s morphological analysis. For the modern sample the species’ name and sample ID are in black font. Posterior probability values of statistical support for each node are indicated in black font. See Figure S6 for a complete view of the modern samples included in the analysis.

The dated BEAST phylogenies recovered the same major clades as the MrBayes analysis lending support to our overall tree topology (Figure S7). In addition to the topology, we were also able to add approximate divergence dates of these main clades as well as the nodes within the clades. Clade 4 (vicugnas) diverges first from the rest of the SACs approximately ~1,01-1,18 Myr. Following the first divergence, Clade 3 forms approximately ~159-201 kyr with the final splint forming Clades 1 and 2 at ~164-198 kyr. Within the clades we uncover a number of different divergence dates between putative domestic and wild mitochondrial haplotypes.

The Bayesian Skyline Plot (BSP) obtained from modern and ancient maternal sequences show an increase in the Ne of the domesticated clades (Clade 1 and 3) around 8,000-6,000 yr B.P. in accordance with the putative initiation of domestication. Clade 2 (guanacos) and Clade 4 (vicuñas) also present evidence of a demographic expansion, however at a much earlier time ~ 50,000 yr BP (Figure 4). Nevertheless, these two clades also show a modern demographic expansion, albeit less pronounced than for Clades 1 and 3, around the domestication time.

**Figure 4.**
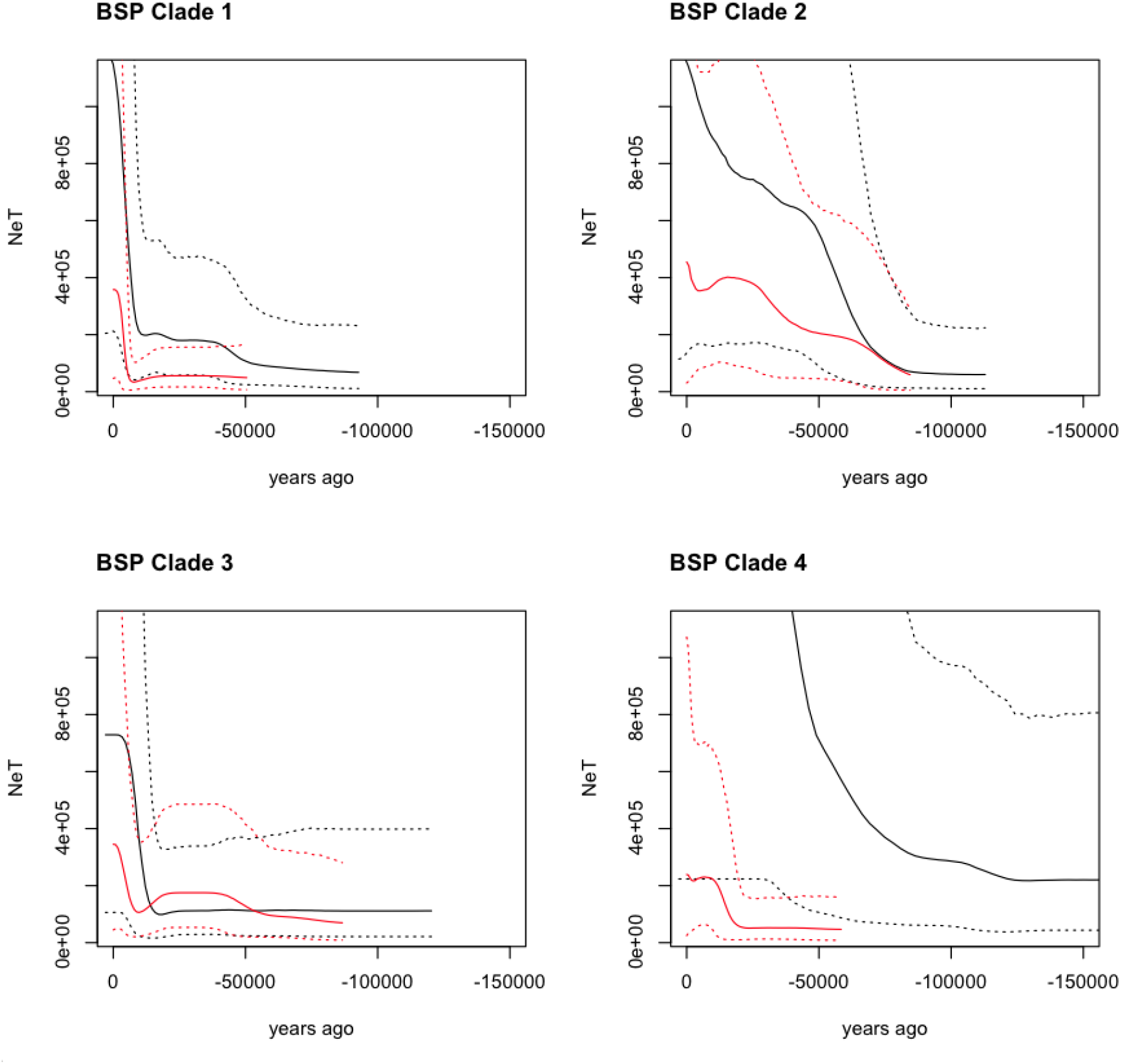
Bayesian Skyline Plot (BSP) derived from the analysis of the ancient and modern camelid data. The x-axis is units of years in the past (from the present - zero, towards (-)150,000 years ago, and the y-axis is equal to NeT (the product of the effective population size and the generation length). The black line is the ancient DNA data set and the red line is the modern DNA data set with the dotted lines indicating the 95% confidence interval of the model and the solid line the mean.

### Control Region Haplotype Network Analysis

To investigate the relationship between the ancient and modern camelids, we analyzed 300 bp of the CR hypervariable I domain. Among the 849 samples analysed, we detected 76 different polymorphic sites and 158 haplotypes divided into two main haplogroups (Figure 5). Thirteen haplotypes were exclusively found in ancient samples, and eleven clustering with the *Lama guanicoe* and domestic camelid haplogroups: *cacsilensis* (66), *cacsilensis-guanicoe* (30), *guanicoe* (76), llamas (25), alpacas (5) (Table S7 and S8).

**Figure 5.**
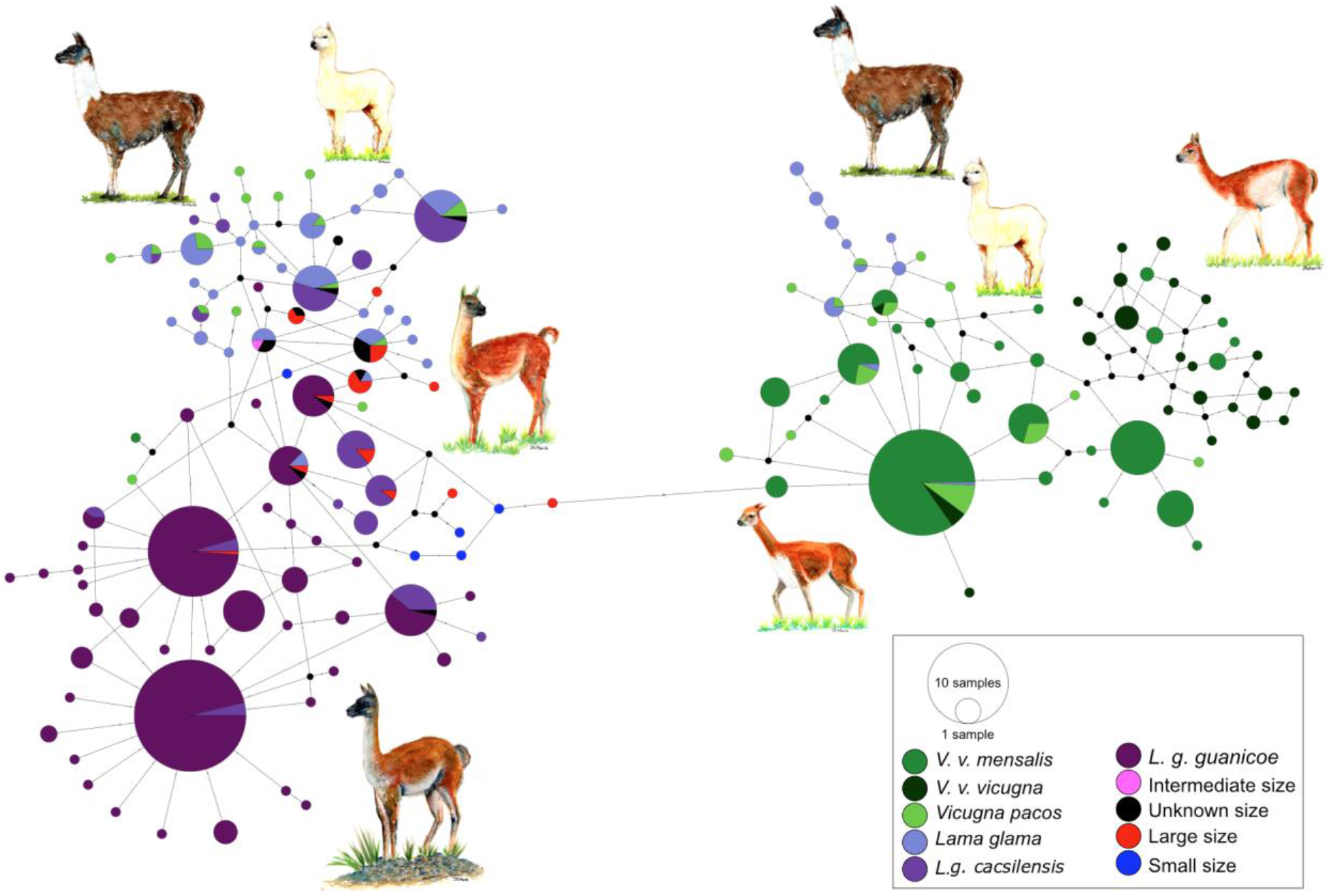
Minimum spanning network of ancient and modern camelid partial control region sequences. A total of 158 different hypervariable I Domain haplotypes found among 849 sequences are shown. Each color represents a camelid subspecies and the ancient sequences. The haplotypes colored in dark purple include the two subspecies of *Lama guanicoe* (*cacsilensis* and *guanicoe*) and light purple represents *Lama glama*. The groups coloured with dark green are formed by the two subspecies of *Vicugna vicugna (vicugna* and *mensalis*), while the light green include the domestic alpacas, *Vicugna pacos*. The aDNA samples from this study are shown in pink, black, red and blue. Each haplotype is represented by a circle with its size proportional to the haplotype’s frequency. Mutations are shown as small perpendicular lines crossing edges connecting haplotypes.

### Temporal network suggests maternal lineage replacement in modern alpacas

A temporal network analysis of the 300 bp of the CR hypervariable I domain was performed using the same dataset as in the PopArt analysis. In total, 849 sequences were included, yielding 158 different haplotypes (42 from ancient sequences and 150 from modern sequences) with only 11 haplotypes being shared by ancient and modern camelids (Figure 6). Despite including a higher number of modern mitochondrial sequences than ancient ones. Most of the shared haplotypes belong to *L. guanicoe guanicoe* and *Lama glama* haplogroups. The temporal network suggests a loss of ancient mitochondrial lineages, mostly in the vicuña.

**Figure 6.**
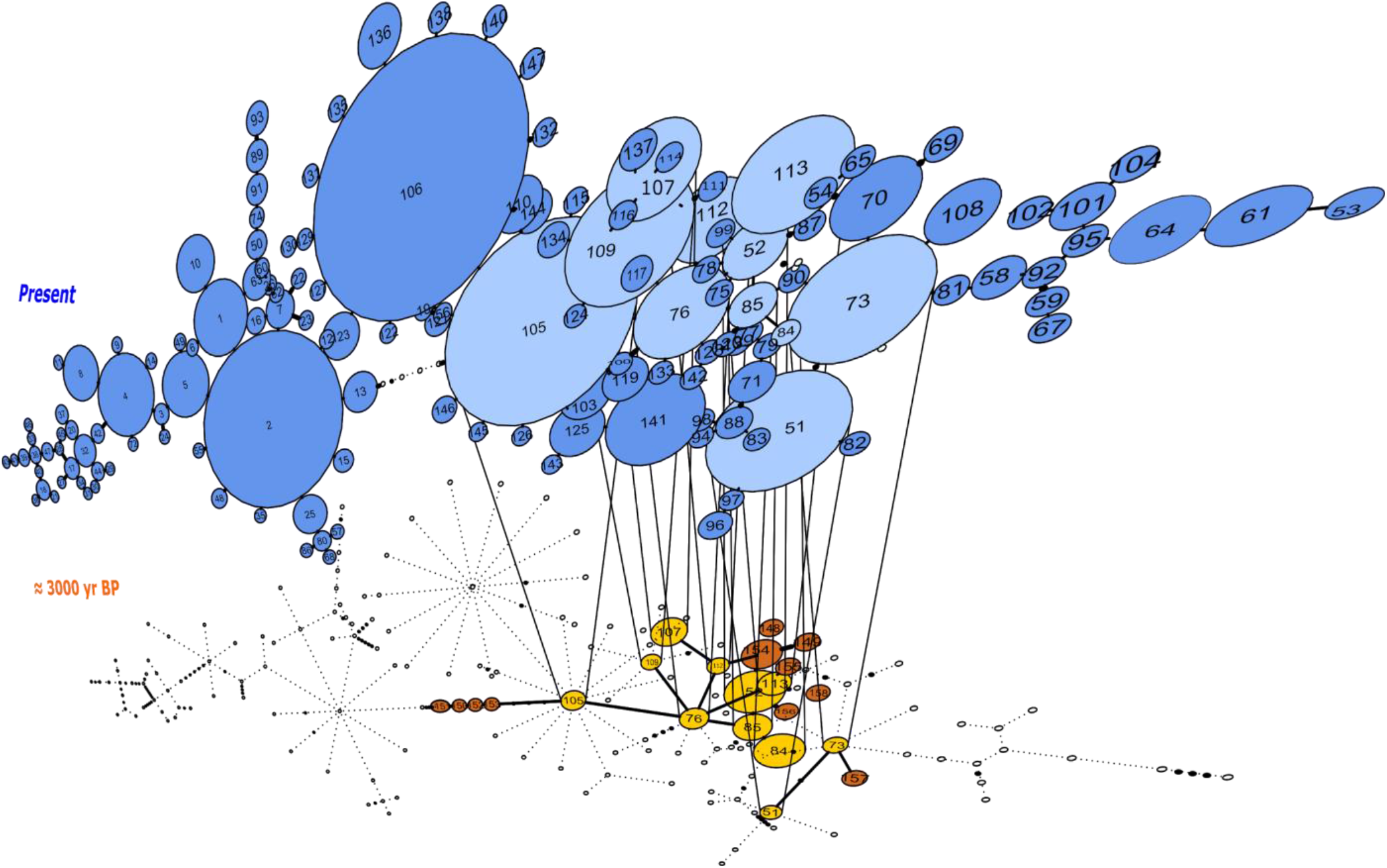
Temporal statistical parsimony haplotype networks for *Lama guanicoe, Lama glama, Vicugna vicugna and Vicugna pacos* with ancient samples. Haplotypes of modern camelid sequences are colored in blue and ancient DNA haplotypes in orange. Each mutation is connected by one black circle. If two haplotypes are separated by one mutation, they are connected by one line. Haplotypes shared by modern and ancient samples are connected by vertical lines showing light blue and yellow color.

Haplotype diversity values were relatively similar between the modern samples (0.9765) and the ancient samples (0.9348). However, given the uneven sample size between modern and ancient samples, we randomly sampled with replacement 10,000 times the modern datasets conditional on a sample size like the ancient samples. For each of the random samples we estimated the haplotype diversity and the number of substitutions observed and used that information to draw a distribution of those two summary statistics of genetic diversity conditional on the ancient DNA sample size. The analysis of all the data (i.e. without differentiating between large and small animals) indicated that the ancient samples do indeed present a high genetic variation given their sample size as their diversity measures fall in the 93^rd^ and 94^th^ percentile of the distribution of the random samples from modern animals (Figure 7 A and D). For the large animals, we observed that the ancient samples fall in the 89^th^ and 95^th^ percentile of the distribution of random samples from the modern *Lama*, while for the small animals, the ancient samples fall in the 98^th^ percentile of the distribution of random samples of *Vicugna* for both statistics.

**Figure 7.**
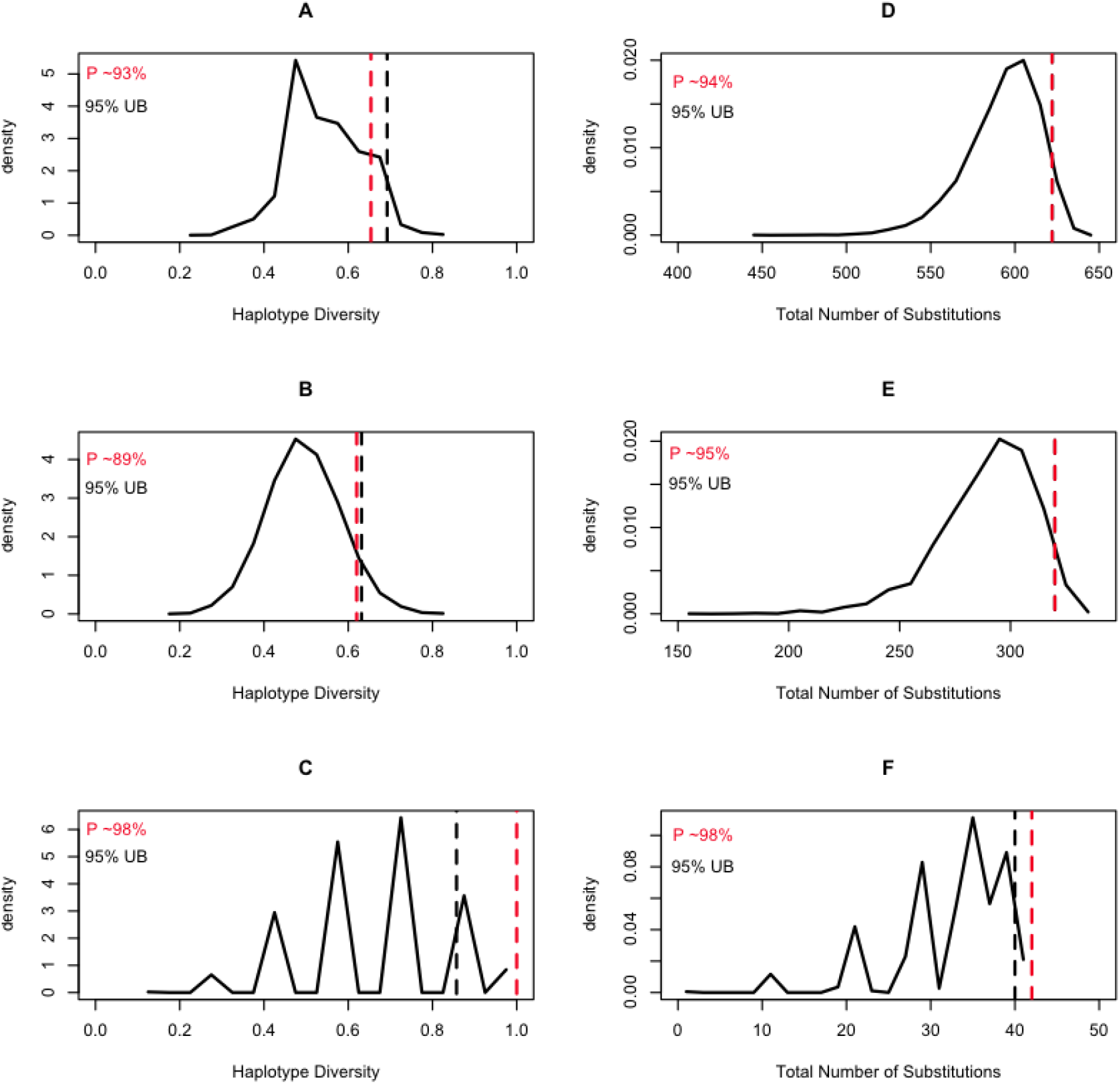
Expected distributions of summaries of genetic diversity conditional on the aDNA sample size. The column on the left shows the results of estimating the haplotype diversity on random samples of the modern animals conditional on the sampling size of the ancient DNA samples. The column on the right shows the results of counting the number of substitutions on the random samples of the modern animals conditional on the sampling size of the ancient DNA samples. The black dashed line is the 95% upper bound of the statistic, and the red dashed line is the position of the ancient DNA data summary statistics of diversity within the context of the random samples.

## Discussion

Here we present to date the most comprehensive dataset of mitogenomes from South American camelids to date, which includes a collection of sequences reflecting archeozoological samples (ancient DNA) from the San Pedro de Atacama desert and dating between ~3,150 and 2,380 years before the present, and a wide collection of modern samples representing the four extant species. We used these data to compare patterns of genetic variation between taxa and across time and to generate insights into hypotheses about the domestication of alpaca and llama. This dataset not only presents unexpected information about the domestication history of South American camelids, it also provides new insights into the loss of genetic diversity in the extant species.

Zooarchaelogy’s foundation is the ability to accurately identify species from animal remains in order to develop an interpretation of human and environmental interactions within an archaeological context **(Peres, 2010)**. While traditionally the field has largely relied on species identification based on comparative analyses with zoological collections or based on researcher’s experience, developments in genetic techniques have facilitated comparisons even when morphological analyses are compromised **(Peres, 2010, LeFebre & Sharpe, 2018)**. Species identification based on bone morphology is not always possible when species are closely related and are only have marginally diverged with respect to each other **(Frantz et al., 2020)**. However, for South American camelids, differentiating between the larger *Lama* animals and the smaller *Vicugna* animals has been shown to work well **(Wheeler 1985, 1995)**. The bone morphological analyses implemented here indicate that the astragalus has a higher discriminatory power to differentiate between larger (*Lama*) animals and the small (*Vicugna*) animals as reflected by the discontinuity in the size measurements between the groups (Figure 2b). While the anterior phalanx has also a good discriminatory power between the two groups there are samples in between them that we could not classify as small or large animals. Nevertheless, the phylogeny generated using 130 ancient and modern camelid mitogenomes places the vast majority of the ancient samples on the expected clades based on bone morphology (Figure 3). Namely, the large size animal bone samples and those of intermediate size were grouped with modern samples of *Lama* (95%) with the exception of two samples that grouped with the *Vicugna vicugna* clade (Clade 4). The converse also occurred with the majority of the small sized animal bone samples that clustered in Clade 4 (80%) with the exception of three (i.e. C69, C77 and C70 from Tulán-54) that grouped with *Lama* (Clades 2 and 3).

### Domestication history of Llamas and Alpacas

Few domestic animals have had their origin as contested like the llamas and alpacas. While their wild relatives, the guanaco and the vicuña, have been determined to have diverged from their common wild ancestor two to three million years ago **(Stanley et al., 1994, Wheeler 1995, Fen et al., 2020)**, the domestic llama and alpaca are assumed to have been domesticated 6,000 to 7,000 years ago **(Wheeler, 1995)**. However, a lack of congruence between methodological analyses have resulted in alternative domestication scenarios being put forward. If the llama was domesticated from guanaco and alpaca from vicuña, we would have expected the phylogenetic tree to group those pairs of taxa accordingly. However, our analyses place the alpaca, with the exception of two samples, within the genus *Lama* which also includes guanaco, modern llamas, and the archaezoological samples of large and intermediate sized animals. While the modern guanacos form their own clade (Clade 2) with one guanaco falling out of that clade and grouping with llamas, they do not form the most closely related clade to the vicuña. Instead, llamas are a paraphyletic group with two main branches (Clades 1 and 3) with the guanacos as sister clade to Clade 1. Furthermore, the alpaca group with llamas in both of their clades supporting the hypothesis that llamas and alpacas were both domesticated from guanacos (hypothesis 3) while vicuña was possibly never domesticated. Such scenario would assume, however, that no alpaca should cluster with vicuñas and that the clade formed by llamas and alpacas has the guanaco as sister taxon **(Renneberg, 2008)**. However, such a partition of samples is not observed in our data. Instead, our mitogenomic phylogenetic tree suggest a modification of such hypothesis where llamas derive from two separate guanaco clades, with the wild component of Clade 3 now extinct, while the extant guanacos are the group from which the other llama clade (Clade 1) derived from. Such a scenario is consistent with the zooarchaeological and genetic evidence for multiple llama domestication centers across the Andes, **(Wheeler et al., 2006; Kadwell et al., 2001; Bruford et al., 2003)** including sites in Northeastern Argentina and Northern Chile **(Mengoni-Goñalons & Yacobaccio, 2003 and 2006; Cartajena, 2009; Núñez & Santoro, 1988; Olivera, 2001; Barreta et al., 2013)** and in the Peruvian central Andes **(Mengoni-Goñalons & Yacobaccio, 2006; Moore, 2011)**.

The domestication hypothesis described above would imply that the alpaca was simultaneously domesticated along with the llama from the two guanaco ancestors based on the clustering of alpacas and llamas in our phylogenetic tree. While that could have happened, it is also possible that alpacas are the outcome of hybridization between llamas and vicuñas (**Wheeler, 1995; Renneberg, 2008**). Alpacas are assumed to have been domesticated independently of llama and probably earlier too **(Wheeler, 2016)** with a Peruvian archaeological time series showing the transition from human hunting of wild vicuñas and guanacos (9,000 - 6,000 years ago) to the establishment of early domestic forms (6,000-5,500 years ago) and the development of human societies centered around pastoralism focused on alpacas and llamas management at high altitude starting 5,500 years ago (**Wheeler, 1986; 1999**). Yet, so far, no evidence of domestication of alpacas has been found outside the domestication centre in central Peru (for which no archaeozoological or modern samples, that could cluster with the vicuña samples, are thus far available). However, the archaeological evidence indicates that precolumbian alpaca populations were substantially larger than today and that a dramatic bottleneck occurred during the Spanish conquest (Wheeler 2016). Thus, it is possible that some of the genetic information about multiple domestication centres for alpaca, as observed in llama, does not exist anymore. Alternatively, the archaeozoological samples of Tulán that are putative alpacas may represent one of those non central Peruvian alpaca domesticated lineages resulting from the cross of female llamas with male vicuñas resulting in phenotypical alpacas that carry a *Lama* mitogenome.

### Early Hybridization of domesticated camelids

Modern alpacas and llamas are the outcome of an extensive hybridization that has been attributed to the species near demise following the Spanish conquest **(Kadwell et al., 2001, Casey et al.,2018, Fen et al., 2020)**. During the Spanish conquest camelid populations were drastically reduced (**Gade, 2013**) probably due to a lack of understanding of the superior quality of their fiber for the textile industry, the interest in the introduction and establishment of cattle and sheep for the production of wool that was in high demand in the Low Countries, and as a measure to control local populations by destabilizing their means of sustenance. Consequently, the paralleled decline in the Indigenous population and the disappearance of their camelid breeding tradition due to the lack of a written system, resulted in a small domestic camelid population that largely was left unmanaged and which could interbreed **(Wheeler et al., 2006, Marin et al., 2007b, Barreta Pinto, 2012)**. Hybridization between llamas and alpacas has become such an extensive issue that up to 40% of llamas and 80% of alpacas show evidence of hybridization (Kadwell et al 2001), and through recent whole genome sequencing efforts, Fen et al. **(2020)** reported that up to 36% of the alpaca genome is likely to be derived from llama hybridization. While the results presented here do not disagree with the hybridization patterns described elsewhere, the observation of hybrid animals among the archaezoological samples (i.e. those samples that present a discordance between their taxon identification based on bone morphology and their molecular taxonomic identification) suggests that the hybridization history of the domestic camelids is yet far from understood. In fact, the hybrid samples (i.e. C69, C77 and C70) present in Tulán (3,400-2,300 yr BP) not only further support the presence of domestic camelids associated to the human settlements in the region, they also provide information about breeding practices by these ancient formative pastoralist communities. Such ancient hybridization is consistent with hypotheses of human controlled crossings between species during the domestication period and afterwards to maintain variation of interest (e.g. adaptive; **Moore, 2011; Marshal et al., 2014)**.

### Loss of genetic variation

The demographic analyses of the ancient and modern samples of each clade in our tree revealed contrasting demographic histories across the dataset. The BSP inferred from the wild animals (Clades 2 and 4) result in substantial demographic expansions that started in the middle of the previous interglacial and continued expanding throughout the last glacial period. Such observation is consistent with an increase in habitable land for *Lama* and *Vicugna* as their distribution ranges probably shifted north and towards lower altitudes as the far South and mountaintops started to freeze. Contrastingly, the ancient domestic samples from Clades 1 and 3 present a roughly stable old demography (Clade 3) or a comparatively moderate population expansion at the onset of the last glacial period (Clade 1). Interestingly, the four clades show a demographic decline starting near ~10,000 yr BP reaching its lowest point around ~6,000 years ago. These dates coincide with the establishment of human populations throughout the upper Andes ~9,000 yr BP **(Aldenderfer, 1999)** and the establishment of a specialized hunting for vicuña and guanaco (9,000-6,000 yr BP) prior to the onset of the domestication process **(Wheeler et al., 1976**). The brief reduction in effective population size is recovered by a population expansion also present in the four clades but exacerbated on the clades with domestic animals where the recovery is substantially larger (Clades 1 and 3; Figure 4). The larger population expansion in Clades 1 and 3, rather than on the clades with wild animals (Clades 2 and 4), is consistent with an increase in domestic animals population size following the onset of the domestication process and the breeding of camelids with traits of interest **(Wheeler, 1995; Thompson et al., 2006; Baied & Wheeler, 1993)**.

The comparative analysis of both the demographic trend observed with the archeozoological samples relative to that observed with the modern samples or between genetic diversity in both datasets, indicates that the ancient camelids presented a much larger genetic pool than the modern ones. In the BSP it can be seen that the plots obtained from the ancient samples (albeit fewer than the modern ones) result in substantially larger effective population sizes and coalescent times (Figure 4), reflecting higher genetic variation in those samples. Similarly, the genetic diversity measured in terms of haplotype sharing between ancient and modern samples, or the number of haplotypes and total number of substitutions observed in both datasets (Figure 7), also show that the ancient samples presented significantly more genetic variation than the modern ones, with the vicuña having lost the most genetic variation across time. The latter is consistent with the evidence for bottlenecks in the species **(Wheeler et al., 1995; Casey et al., 2018; González et al., 2019; Fen et al 2020)**.

In this study, we presented new evidence for the domestication history of South American camelids and propose a model of domestication that includes an ancient guanaco population that no longer exists. Furthermore, we propose that the alpaca may have a more complex evolutionary history, probably deriving from the guanaco or at least including an important component of that species genetic variation in the alpaca genetic make-up, as well as hybridization between these species having played a role since their domestication time. Lastly, through dramatic changes in their history, the four camelid species show evidence of having lost a very large proportion of their genetic variation with modern populations only representing a small fraction of that. While some of the extant camelid populations have been brought back from the brink of extinction in the last 50 years (e.g. Peruvian vicuña), current climate change exacerbating the aridification of the upper Andes, land conversion for agriculture and a general lack of appetite from local governments to invest in these native species of agricultural value, are likely to drive these camelids into further trouble.

