## Supplementary material for "Ancient DNA Reveals the Lost Domestication History of South American Camelids in Northern Chile and Across the Andes"

### Material and Methods

*Biogeographical area*

### Description of the fossil site and sampling Archaeological sites: Tulán-54 & Tulán-85

The geographic area is located in the II Region of Chile, Antogasta, specifically in the western slope of the Puna de Atacama (22°-24°S) **(Figure S1).** Situated along the Tulán transect, which stretches over 15km southeast of the Salar de Atacama basin, are placed the two archaeological sites for this study. During the Early Formative (ca. 5000 yr BP), groups developed a mixed economy sustained by hunting, gathering and camelid breeding while integrating an important ritual component, new technologies and the development of ceremonial and architecture. This period (c. 3360–2370 yr BP) is well represented by two important settlements (Tulán-54 and Tulán-85). *Tulán-85*, located on the border of the resource-rich salt flat, contained few residential structures but comprised an extensive and deep refuse mound resulting mainly from production activities (Núñez 1992) **(Figure S3c)**. *Tulán-54 site*, located in the upper part of the ravine (3000 masl), represents a ceremonial center with monumental architecture and burials where the repetition of ritual events is expressed throughout the remains of which filled the temple in a span of 200years **(Figure S2, S3a, S3b).** Tulán-54 site emerges as a religious axis within a context of macroregional networks. Human groups built this particular temple oriented towards the social cohesion of groups with a wide mobility between the Tulán ravine and other settlements from the Puna (Núñez et al., 2017). Tulán-54 and Tulán-85 have been previously associated with an early herding and hunting economy. Human populations started since 3000 yr BP ago to choose wild camelids to have their own herd so the identification of domesticated camelids in these sites could be reasonable.

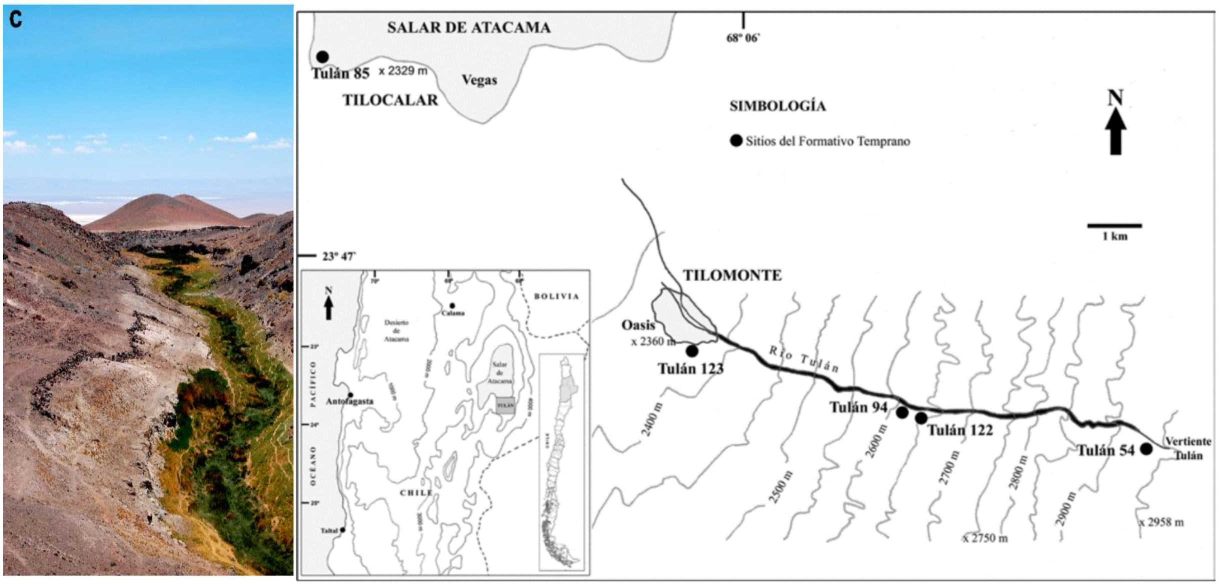


**Figure S1.Location of sites assigned to the Early Formative Period in the *Tulán* ravine (Figure taken from *López* et al. 2013).**

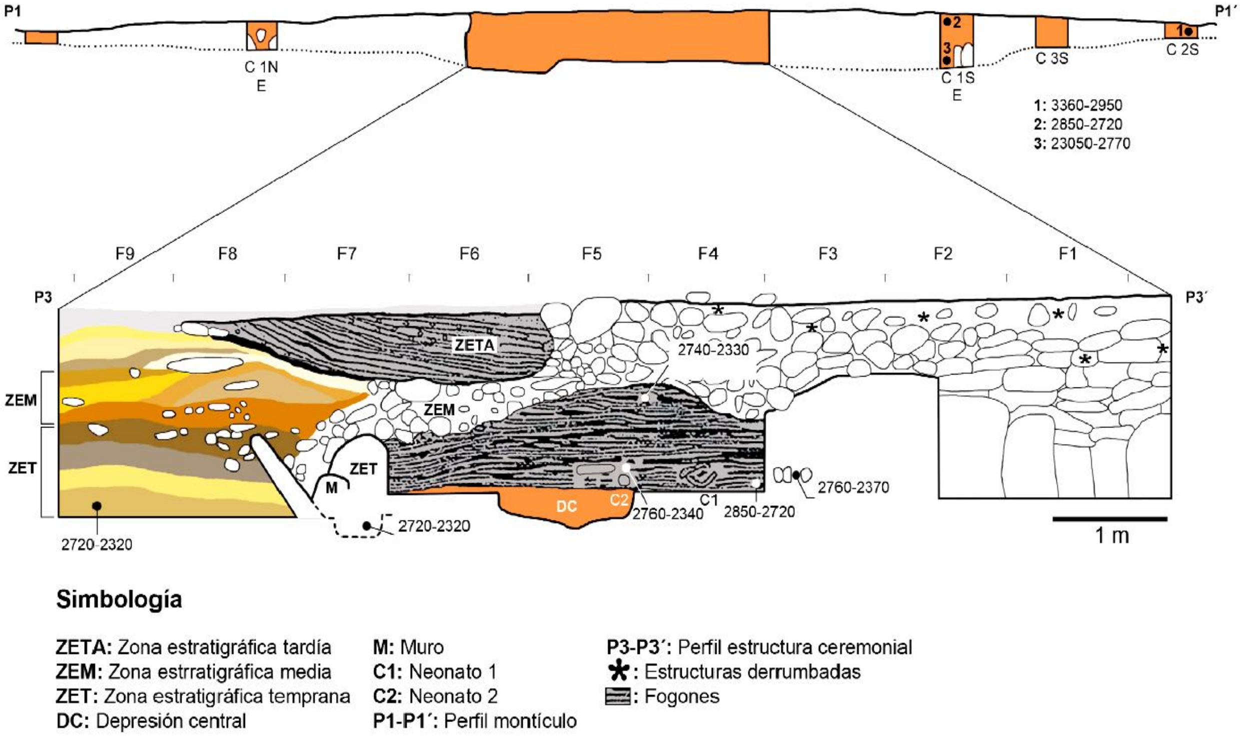


**Figure S2. Transversal stratigraphic profile of the Tulán-54 site, showing details of the deposits inside the temple. Picture taken from Núñez et al., 2017.**

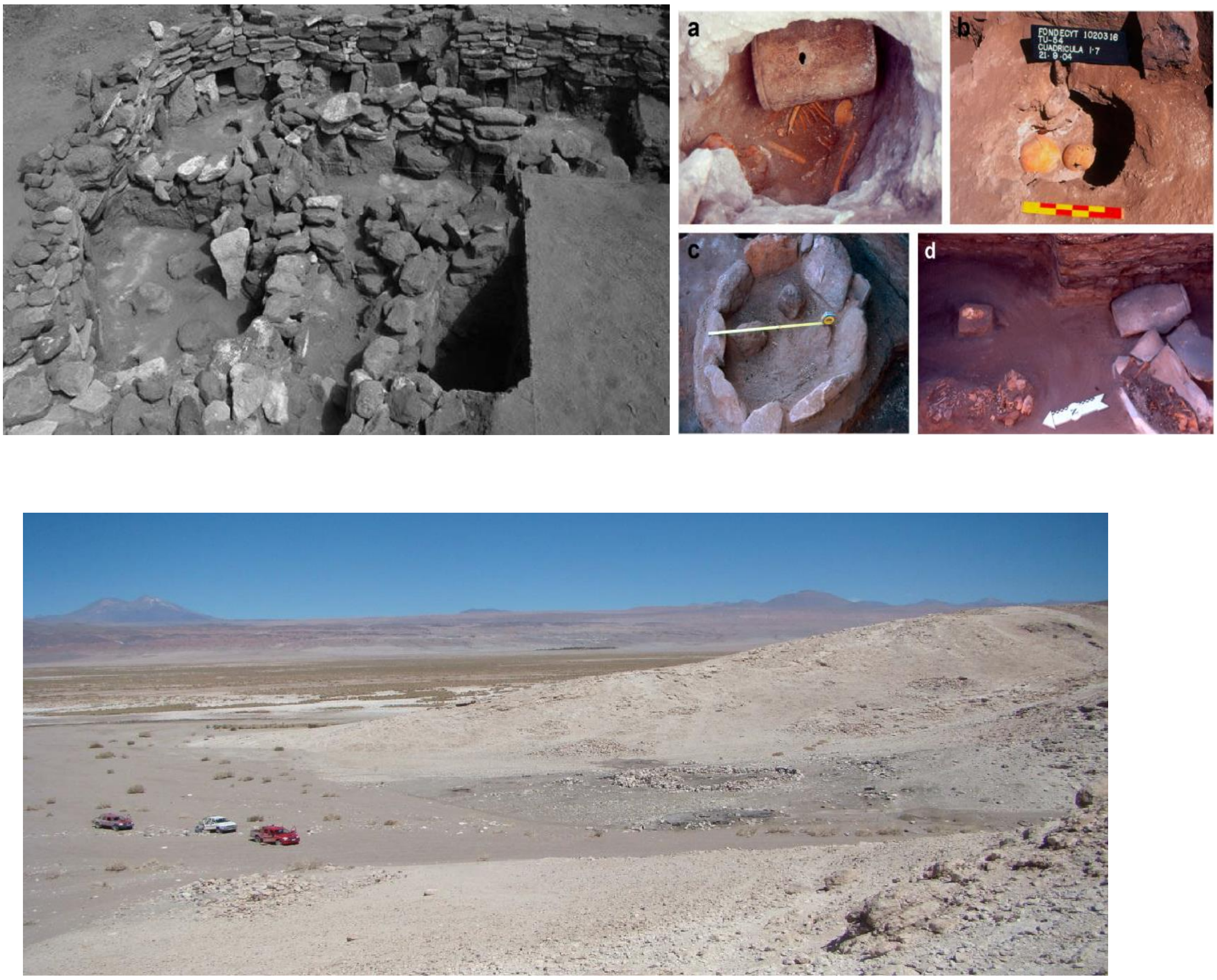


**Figure S3.a) Partial view of the ceremonial structure at Tulán-54. b) Human burial with offering inside the ceremonial site at Tulán-54. c) Panoramic view of Tulán-85.**

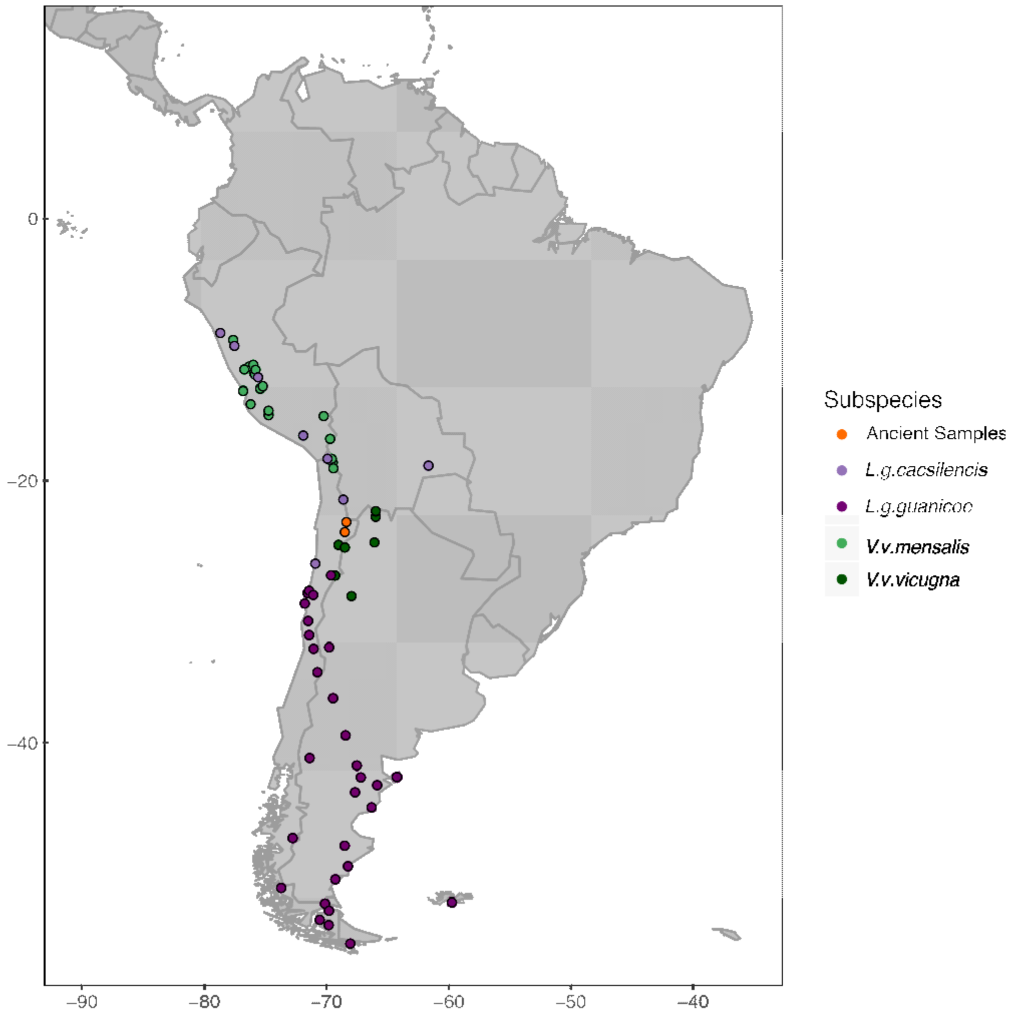


**Figure S4. Distribution map of modern camelid samples included in the Bayesian phylogenetic analysis. Each colour represent one subspecie of camelid. Orange spots belongs to the location of the archaeological sites, Tulán-54 and Tulán-85. Data collected from Dr. Juan Carlos Marín.**

*Map Damage*

**Figure S5. DNA fragmentation and nucleotide mis-incorporation plots for samples from Tulán54, Tulán85, Tulán-52 and Tulán94.**

DNA fragmentation patterns are shown for the 10bp preceding reads starts (positions -1 to -10 on the left panels of each base composition profile) and the 10bp following read ends (post-adapter and/or quality trimming; positions 1 to 10 on the right panel of each base composition profile= Nucleotide mis-incorporation patterns (3$^{rd}$ row) are shown for the first 5 nucleotides sequenced and the last 50 (post-adapter and/or quality trimming), with C ---> T substitution frequencies shown in red, and G--->A substitution frequencies shown in blue.

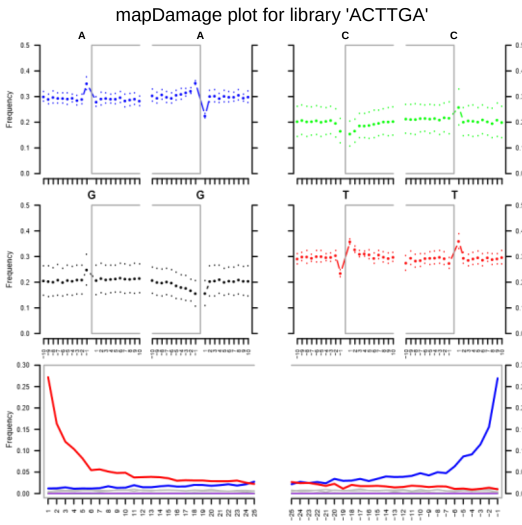


**TU-54: C55** ***Vicugna pacos clade***

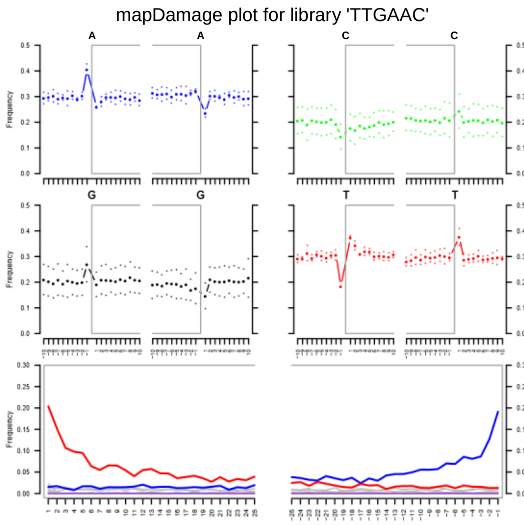


**TU-85: C36** ***Lama guanicoe*** **clade**

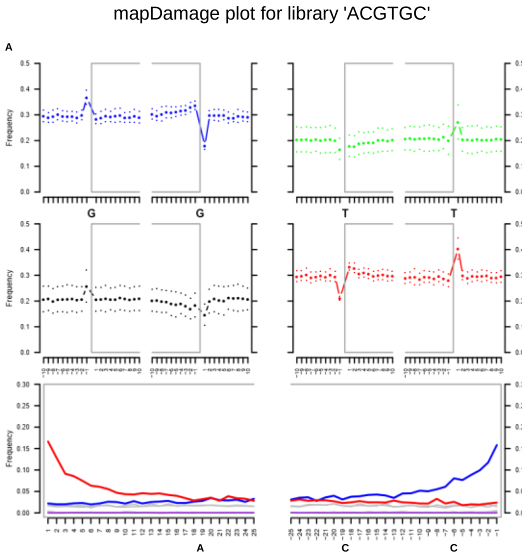


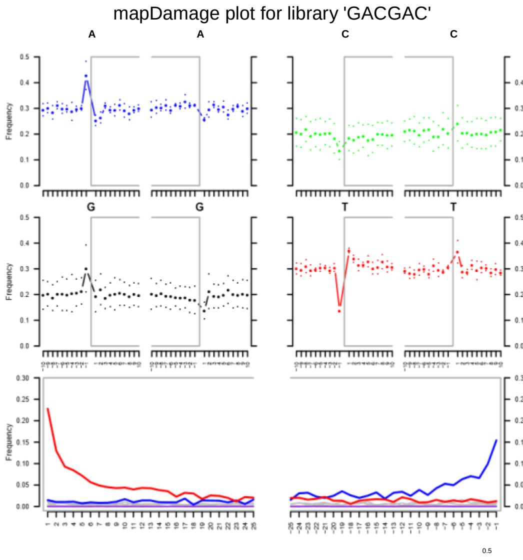

**TU-54: C34** *Vicugna vicugna* clade

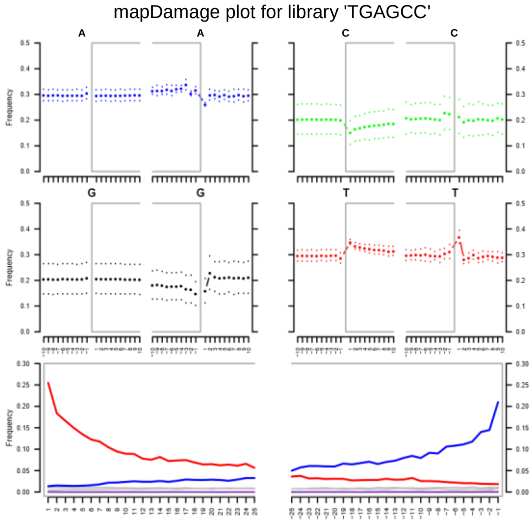


**TU-52: C46** *Vicugna pacos* clade

**TU-85:C6** *Lama glama* clade

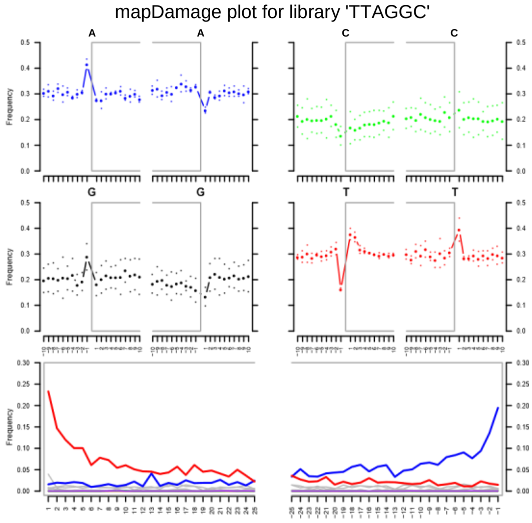


**TU-94: C52**

**Figure S6. Mitogenomic phylogenetic tree.** Bayesian reconstruction of phylogenetic relationships between ancient and modern camelids using complete mitochondrial genome data. All names of the modern samples used in this analyses are included in the phylogenetic tree.

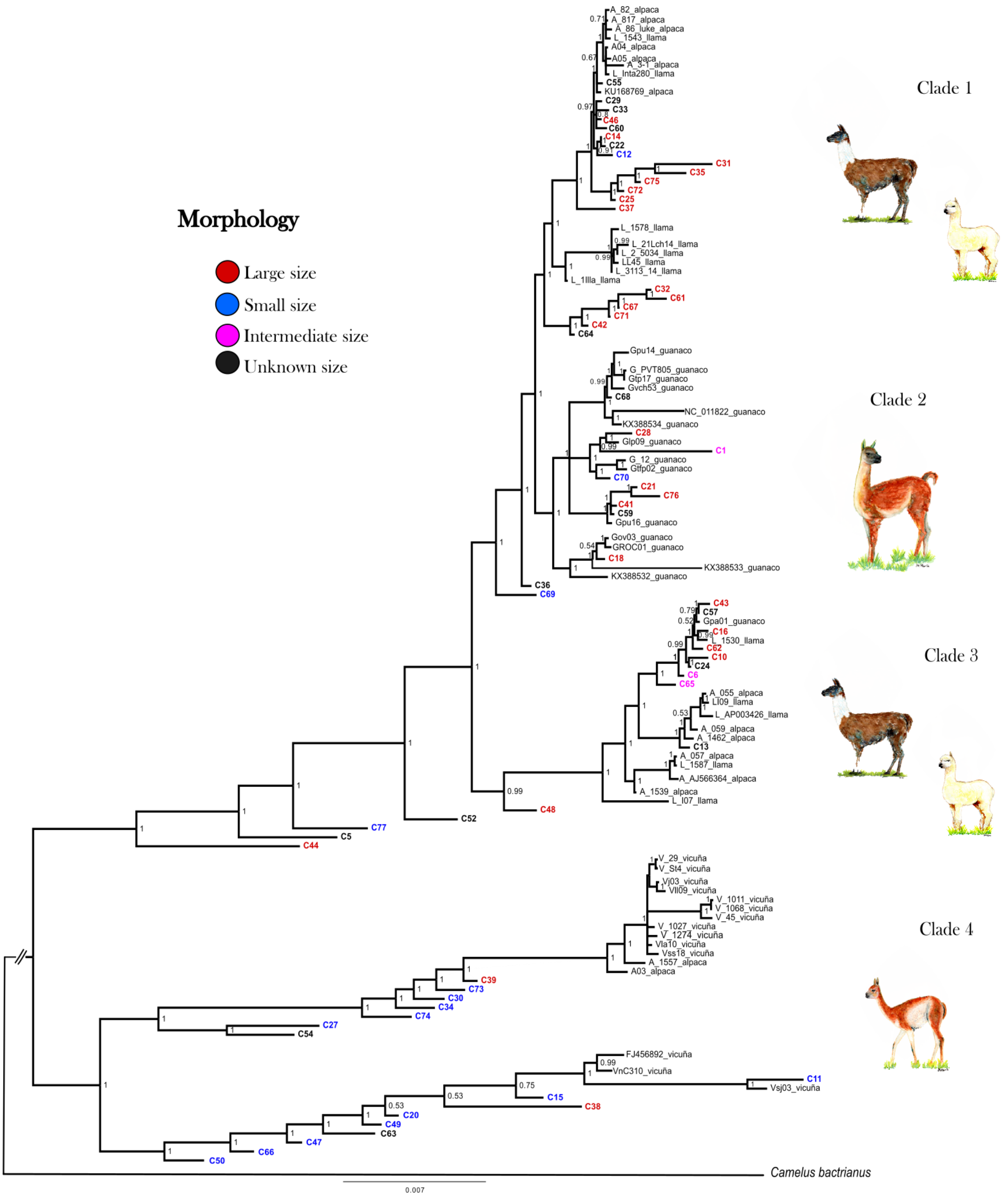

**Figure S7. Dated mitogenomic phylogenetic tree. Phylogeny and molecular timescale of South American camelids including Old World camels as an outgroup.** Values in brackets refer to the gap of time when every clade diverge from another. A) Enlargement of the red clade B) Enlargement of the yellow clade C) Enlargement of purple clade.

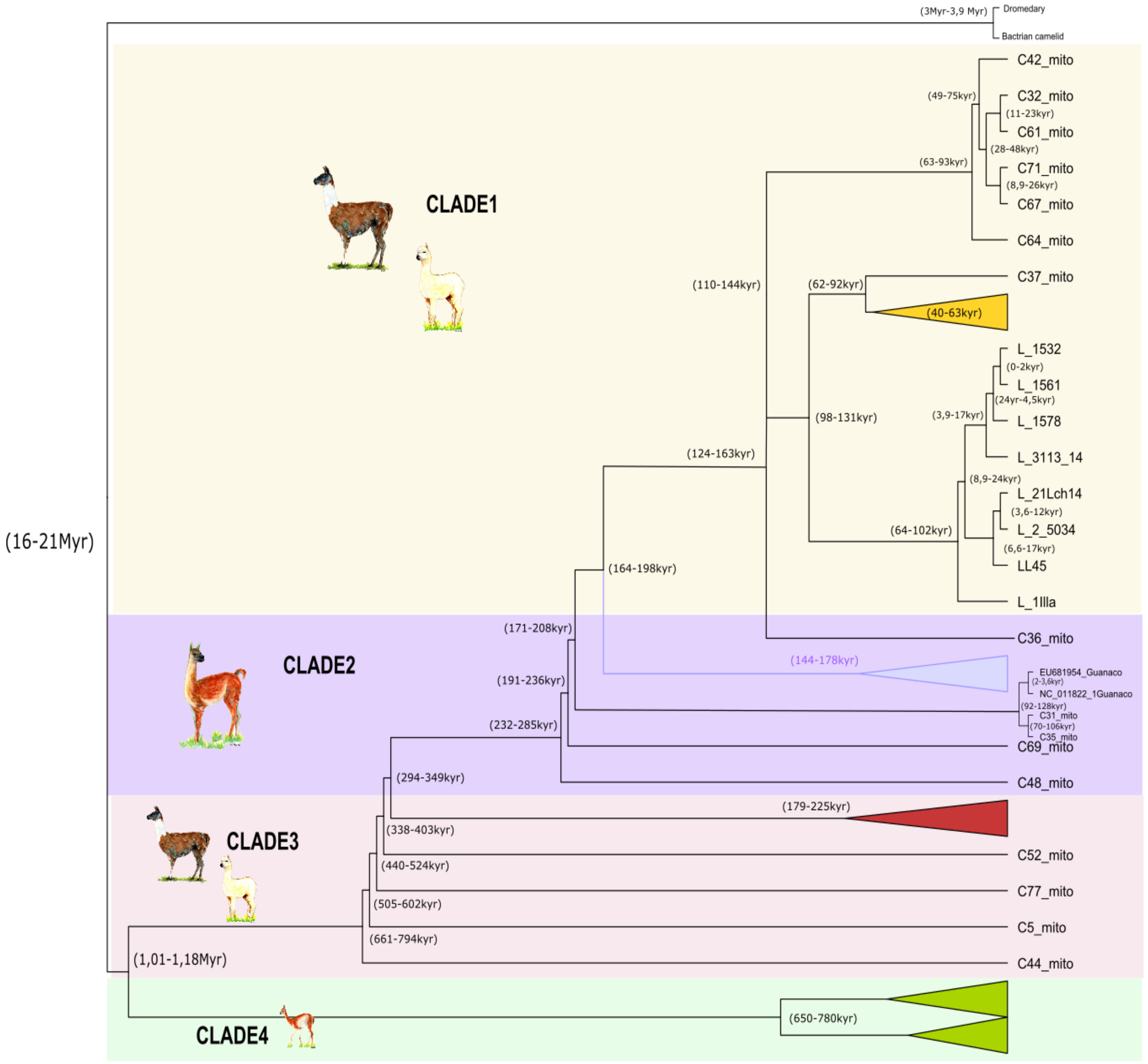

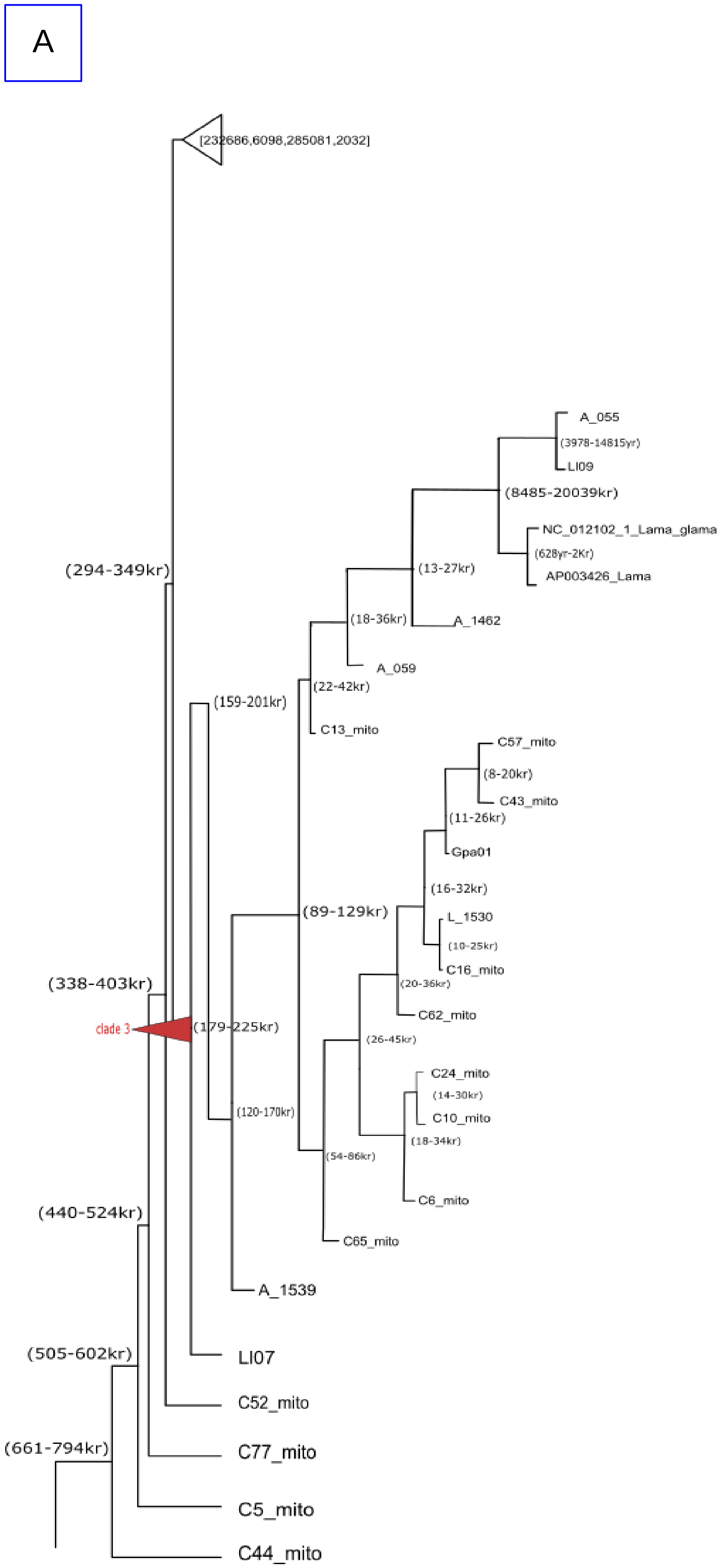

A
[232686,6098,285081,2032]
A_055
(3978-14815yr)
LI09
(8485-20039kr)
NC_012102_1_Lama_glama
(628yr-2Kr)
AP003426_Lama
(294-349kr)
(13-27kr)
(18-36kr)
A_1462
A_059
(159-201kr)
(22-42kr)
C13_mito
C57_mito
(8-20kr)
C43_mito
(11-26kr)
Gpa01
(16-32kr)
(89-129kr)
L_1530
(10-25kr)
C16_mito
(338-403kr)
(20-36kr)
C62_mito
clade 3
(179-225kr)
(26-45kr)
C24_mito
(14-30kr)
C10_mito
(120-170kr)
(18-34kr)
(54-86kr)
C6_mito
(440-524kr)
C65_mito
A_1539
(505-602kr)
LI07
C52_mito
(661-794kr)
C77_mito
C5_mito
C44_mito

B

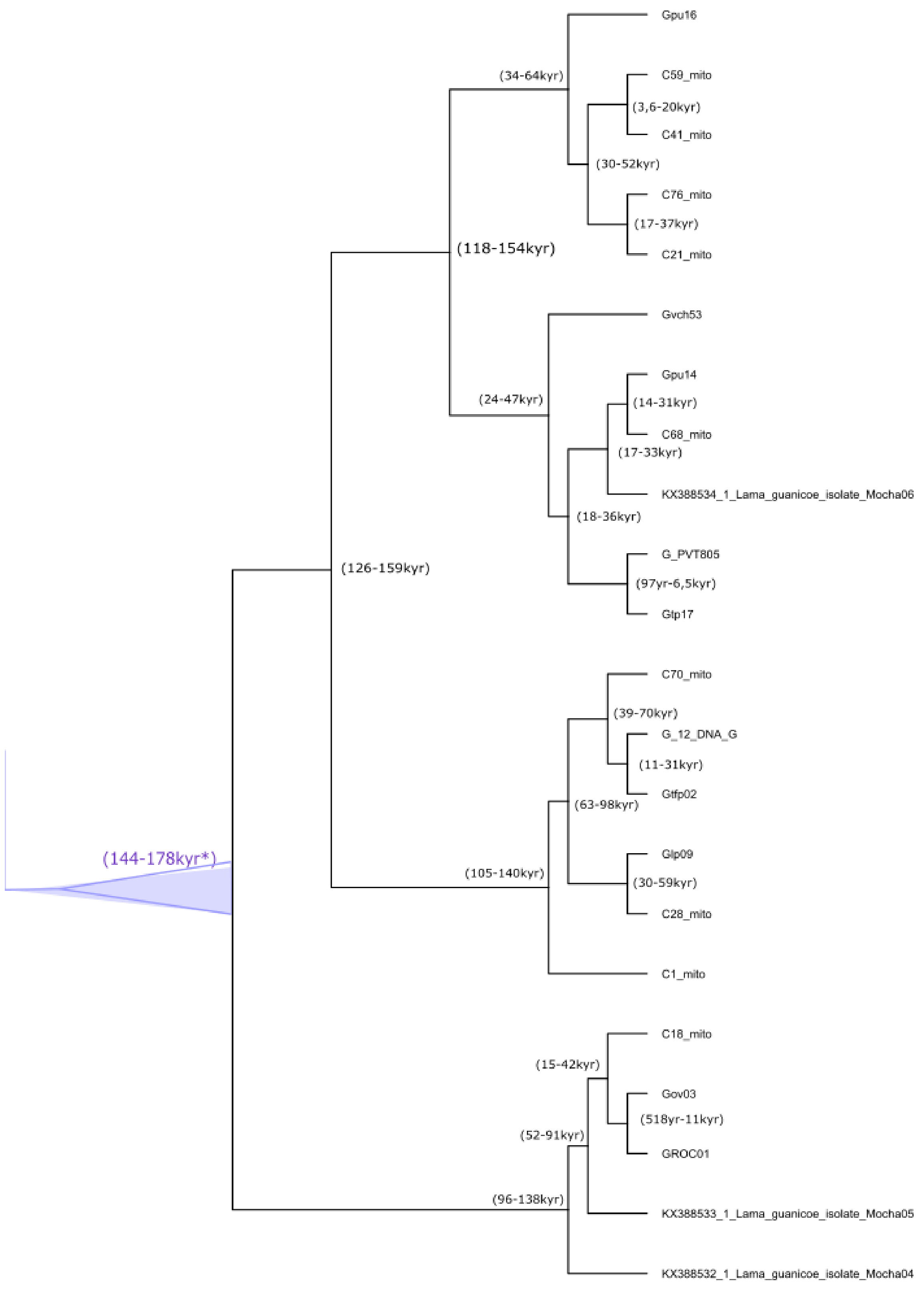
Gpu16
(34-64kyr)
C59_mito
(3,6-20kyr)
C41_mito
(30-52kyr)
C76_mito
(17-37kyr)
C21_mito
(118-154kyr)
Gvch53
Gpu14
(24-47kyr)
(14-31kyr)
C68_mito
(17-33kyr)
KX388534_1_Lama_guanicoe_isolate_Mocha06
(18-36kyr)
G_PVT805
(126-159kyr)
(97yr-6,5kyr)
Glp17
C70_mito
(39-70kyr)
G_12_DNA_G
(11-31kyr)
Gtfp02
(63-98kyr)
(144-178kyr*)
Glp09
(30-59kyr)
(105-140kyr)
C28_mito
C1_mito
C18_mito
(15-42kyr)
Gov03
(518yr-11kyr)
(52-91kyr)
GROC01
(96-138kyr)
KX388533_1_Lama_guanicoe_isolate_Mocha05
KX388532_1_Lama_guanicoe_isolate_Mocha04

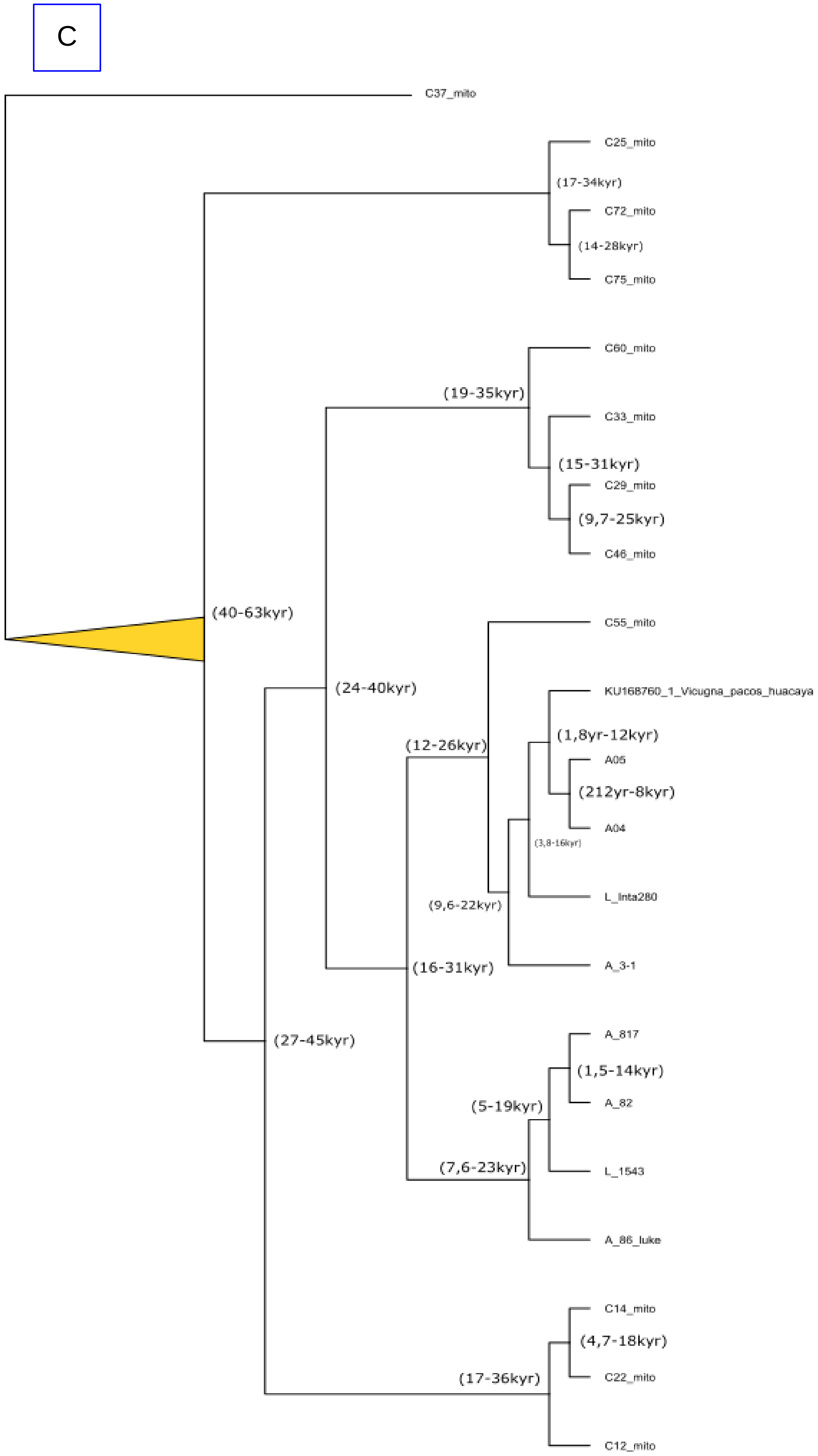
C
C37_mito
C25_mito
(17-34kyr)
C72_mito
(14-28kyr)
C75_mito
C60_mito
(19-35kyr)
C33_mito
(15-31kyr)
C29_mito
(9,7-25kyr)
C46_mito
(40-63kyr)
C55_mito
(24-40kyr)
KU168760_1_Vicugna_pacos_huacaya
(1,8yr-12kyr)
(12-26kyr)
A05
(212yr-8kyr)
A04
(3,8-16kyr)
L_Inta280
(9,6-22kyr)
A_3-1
(16-31kyr)
(27-45kyr)
A_817
(1,5-14kyr)
A_82
(5-19kyr)
L_1543
(7,6-23kyr)
A_86_luke
C14_mito
(4,7-18kyr)
C22_mito
(17-36kyr)
C12_mito
